# An oncogenic isoform of septin 9 promotes the formation of juxtanuclear invadopodia by reducing nuclear deformability

**DOI:** 10.1101/2023.06.18.545473

**Authors:** Joshua Okletey, Dimitrios Angelis, Tia M. Jones, Cristina Montagna, Elias T. Spiliotis

## Abstract

Invadopodia are extracellular matrix (ECM) degrading structures, which promote cancer cell invasion. The nucleus is increasingly viewed as a mechanosensory organelle that determines migratory strategies. However, how the nucleus crosstalks with invadopodia is little known. Here, we report that the oncogenic septin 9 isoform 1 (SEPT9_i1) is a component of breast cancer invadopodia. SEPT9_i1 depletion diminishes invadopodia formation and the clustering of invadopodia precursor components TKS5 and cortactin. This phenotype is characterized by deformed nuclei, and nuclear envelopes with folds and grooves. We show that SEPT9_i1 localizes to the nuclear envelope and juxtanuclear invadopodia. Moreover, exogenous lamin A rescues nuclear morphology and juxtanuclear TKS5 clusters. Importantly, SEPT9_i1 is required for the amplification of juxtanuclear invadopodia, which is induced by the epidermal growth factor. We posit that nuclei of low deformability favor the formation of juxtanuclear invadopodia in a SEPT9_i1-dependent manner, which functions as a tunable mechanism for overcoming ECM impenetrability.

**Highlights:**

- The oncogenic SEPT9_i1 is enriched in breast cancer invadopodia in 2D and 3D ECM
- SEPT9_i1 promotes invadopodia precursor clustering and invadopodia elongation
- SEPT9_i1 localizes to the nuclear envelope and reduces nuclear deformability
- SEPT9_i1 is required for EGF-induced amplification of juxtanuclear invadopodia

**eTOC Blurb:** Invadopodia promote the invasion of metastatic cancers. The nucleus is a mechanosensory organelle that determines migratory strategies, but how it crosstalks with invadopodia is unknown. Okletey et al show that the oncogenic isoform SEPT9_i1 promotes nuclear envelope stability and the formation of invadopodia at juxtanuclear areas of the plasma membrane.

## INTRODUCTION

Metastatic cancer cells breach tissue barriers by forming specialized structures termed invadopodia ^1–3^. Invadopodia are elongated finger-like protrusions that secrete matrix metalloproteinases (MMPs), which cleave the extracellular matrix (ECM) ^4, 5^. Invadopodia formation is induced by growth factors such as the epidermal growth factor (EGF) or transforming growth factor β (TGFβ), which promote tumor migration and invasion by activating non-receptor tyrosine kinases and the phosphoinositide 3-kinase (PI3K)/Akt signaling pathway ^4, 6–8^. Nucleation of invadopodia occurs at plasma membrane domains with clusters of invadopodia precursor complexes that consist of actin binding proteins (cortactin, cofilin, NCK1) and nucleation-promoting factors (ARP2/3, N-WASP) ^9–12^. Invadopodia precursors mature upon phosphorylation and recruitment of the scaffold protein TKS5, which interacts with phosphatidylinositol 3,4-bisposphate ^11, 13–15^. Concomitantly, actin polymerizes into filaments, which form the structural core of elongating invadopodia ^1^.

The nucleus is increasingly viewed as a mechanosensory organelle with critical roles in the mechanisms and modes of cell migration ^16–18^. Recent studies indicate that ECM degradation by invadopodia is an adaptive response to high nuclear confinement, which results from low deformability of the nucleus and/or an impenetrable ECM of high rigidity and small pore size ^19–22^. Invadopodia often form underneath the nucleus ^23^. The apical tips of invadopodia terminate to nuclear indentations, suggesting that invadopodia physically contact and push against the nuclear envelope ^23^. Invadopodia, though, also exert outward forces that repel and reorganize the ECM around the nucleus ^24^. Additional evidence for a crosstalk between the nucleus and invadopodia comes from the localization of MMP-containing endolysosomes, which position anteriorly to the migrating nucleus in a manner that depends on the linker of nucleoskeleton and cytoskeleton complex (LINC) ^21^. In spite of these findings, whether and how invadopodia crosstalk with the nucleus is little understood.

Septins are a family of GTP-binding proteins, which are structurally and evolutionarily related to the Ras family of GTPases ^25^. *SEPT9* is a septin gene which is amplified in murine models of breast cancer and human breast cancers ^26–29^. *SEPT9* encodes multiple isoforms, and isoform 1 (SEPT9_i1) is the longest isoform with oncogenic properties ^29, 30^. SEPT9_i1 promotes cell proliferation and angiogenesis through interactions with the c-Jun-N-terminal-kinase (JNK) and the hypoxia inducible factor 1a (HIF1a) ^31, 32^. Notably, SEPT9_i1 over-expression enhances cell migration and invasion, as well as ECM degradation in 2D and 3D in vitro assays ^27, 33–36^. Moreover, SEPT9_i1 over-expression and knock-down enhances and impedes, respectively, tumor growth and metastasis in mice ^35, 37, 38^. In metastatic tumor cell lines, SEPT9 mRNA is enriched in pseudopods ^39^, and SEPT2 promotes the maturation of podosomes in endothelial cells ^40^. It is unknown, however, whether septins have roles in the formation and degradative functions of invadopodia.

Here, we sought to investigate if the oncogenic SEPT9_i1 is involved in the formation of invadopodia. Surprisingly, we found that SEPT9_i1 is a component of both invadopodia and the nuclear envelope, and is required for the preferential formation of invadopodia at juxtanuclear sites of the ventral plasma membrane. We propose that SEPT9_i1 amplification enhances cancer cell invasion by promoting the formation of invadopodia at juxtanuclear sites of the plasma membrane, which facilitates the translocation of cancer nuclei through ECMs of poor penetrability.

## RESULTS

### SEPT9_i1 localizes to the invadopodia of breast cancer cells in 2D and 3D ECMs

In metastatic human breast cancer MDA-MB-231 cells, which contain a high copy number of the *SEPT9* gene ^27^, we examined whether the oncogenic isoform SEPT9_i1 is involved in invadopodia formation. We used 2D and 3D assays of ECM degradation to visualize invadopodia in MDA-MB-231 cells, which were treated with EGF to upregulate invadopodia. We first probed for the localization of SEPT9_i1 with an isoform-specific antibody in cells, which were plated on a 2D matrix of fluorescent gelatin. SEPT9_i1 was enriched in broader areas of gelatin degradation that overlapped with the nucleus (Figure 1A) and often localized to perinuclear regions of gelatin degradation (Figure 1B). We observed SEPT9_i1 filaments transverse spots that lacked gelatin (Figure 1B, region 1). Additionally, SEPT9_i1 surrounded and partly filled foci of gelatin depletion (Figure 1B, regions 2 and 3). Fluorescence quantifications showed that SEPT9_i1 is more enriched in smaller areas (< 4 μm^2^) of gelatin degradation (Figure 1C), which suggested that SEPT9_i1 may function in early stages of invadopodia formation and ECM degradation.

**Figure 1.**
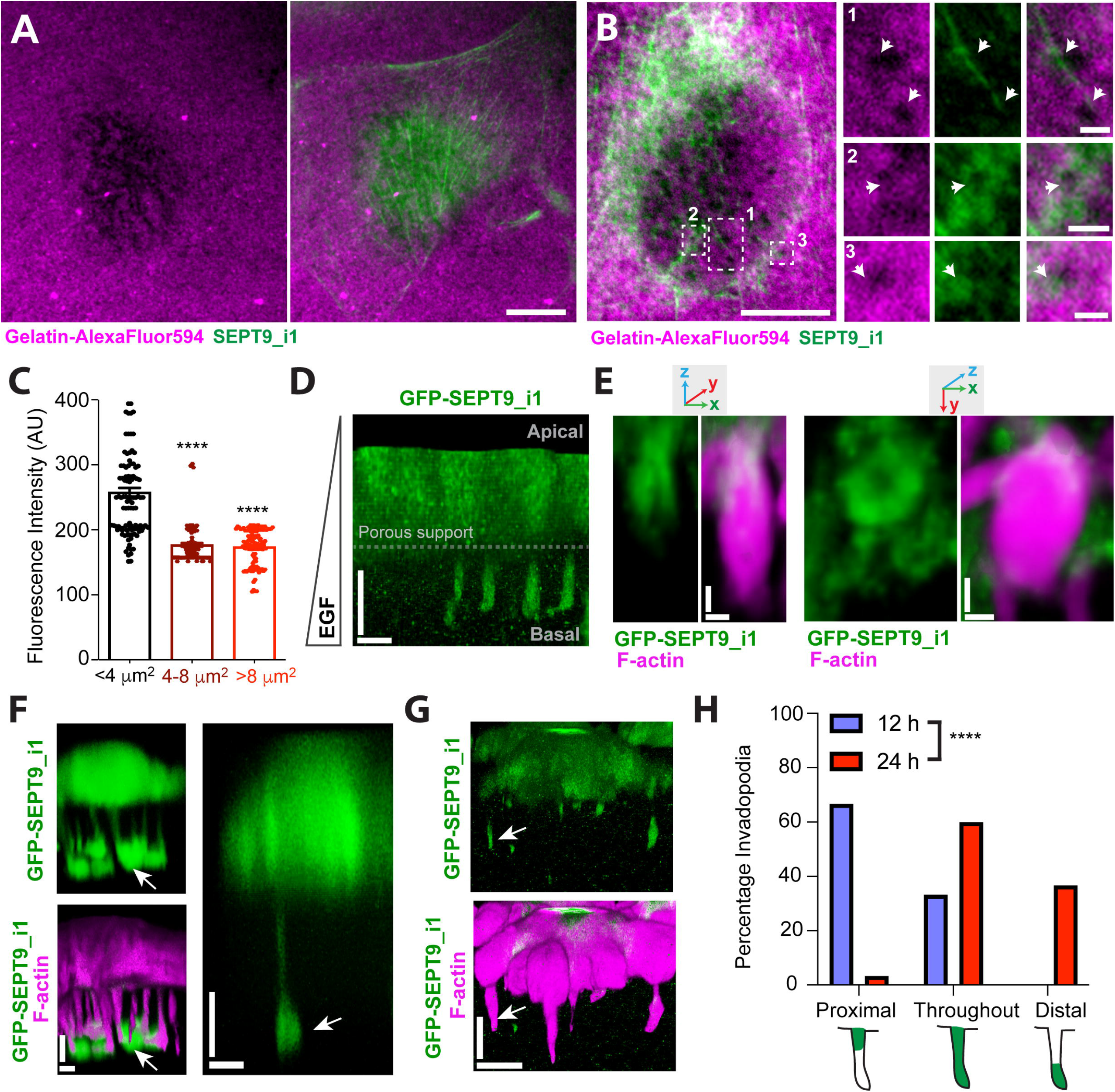
SEPT9_i1 localizes to sites of invadopodia formation and ECM degradation in 2D and 3D. (A-B) Images show MDA-MB-231 cells, which were plated on gelatin-AlexaFluor594 (magenta) for 5 h in EGF (10 nM, 4h) and stained with an isoform-specific antibody against SEPT9_i1 (green). Images show SEPT9_i1 localization to nuclear (A) and perinuclear (B) regions of gelatin-AlexaFluor594 degradation. Three regions of gelatin-AlexaFluor594 degradation are shown in higher magnification (B). Arrows point to a SEPT9_i1 filament traversing two spots of gelatin degradation (1) and SEPT9_i1 accumulations around (2) and inside areas of degraded gelatin (3). Scale bars, 10 μm and 1 μm (insets). (C) Bar graph shows the mean (±SEM) intensity of endogenous SEPT9_i1 fluorescence in surface areas (n = 112) of gelatin-AlexaFluor594 degradation with sizes of <4 μm^2^, 4-8 μm^2^ and >8 μm^2^. Data were analyzed with a non-parametric one-way ANOVA Kruskal-Wallis test with Dunn’s post-hoc analysis for pairwise comparisons. ****, p < 0.0001 (D-E) MDA-MB-231 cells that stably express GFP-SEPT9_i1 were plated on permeable Matrigel- and laminin-coated Transwell membranes for 12 h in the presence of an EGF gradient, which induced invadopodia extension toward the basal compartment. Images of 3D-rendered confocal z-stacks show a side-view of GFP-SEPT9_i1 enrichment in the shaft of invadopodia (D) and at the base of invadopodia, where GFP-SEPT9_i1 has a ring-like appearance (E). Scale bars, 10 μm (D) and 2 μm (E). (F-G) MDA-MB-231 cells that stably express GFP-SEPT9_i1 were plated on permeable Matrigel- and laminin-coated Transwell membranes for 24 h in the presence of apicobasal EGF gradient. Images show side views of confocal z-stacks after 3D-rendering. GFP-SEPT9_i1 localizes throughout the shaft and tip of invadopodia (F), or is enriched in distal segments of invadopodia (G). Arrows point to GFP-SEPT9_i1 to the distal tips of invadopodia. Scale bars, 5 μm (F) and 10 μm (G). (H) Quantification of GFP-SEPT9_i1 localization in the 3D invadopodia of MDA-MB-231 cells, which were plated on Matrigel-/laminin-coated Transwell membranes for 12 or 24 h in the presence of an EGF gradient. Bar graph shows percentage of invadopodia (n = 30) with GFP-SEPT9_i1 localizing to the base of invadopodia (proximal), along the shaft and tip of invadopodia (throughout), and the distal end of invadopodia (distal). Data were analyzed with the chi square test. ****, p < 0.0001

Super-resolution confocal microscopy showed that SEPT9_i1 filaments span the ventral surface of the nucleus and colocalize with actin filaments in cells adhered on gelatin (Figure S1A) or collagen (Figure S1B). A few SEPT9_i1 filaments, however, did not overlap with actin (Figure S1A-B, arrowheads), and SEPT9_i1 was not enriched in ventral actin rosettes and lamellipodia (Figure S1A-B). SEPT9_i1 colocalized with SEPT7 and SEPT2 (Figure S1A-D), indicating that SEPT9_i1 was organized in complexes consisting of the canonical SEPT2/6/7/9 protomer as previously reported ^41, 42^. SEPT9_i1 was also present at regions of the nuclear rim as shorter filaments that ran along (Figure S1C, i) or intersected orthogonally the nuclear envelope (Figure S1E). In apical sections of MDA-MB-231 cells, SEPT9_i1 colocalized with actin filaments that stretched along the dorsal surface of the nucleus resembling the transmembrane actin-associated nuclear (TAN) lines (Figure S1F) ^43^.

To rule out the possibility that SEPT9_i1 localization to sites of gelatin degradation was an artifact of antibody staining and/or 2D culture, we induced invadopodia formation in a 3D chemoinvasion assay using MDA-MB-231 cells that were stably transfected with GFP-SEPT9_i1. Cells were plated for 12 or 24 hours on Matrigel/laminin-coated transwell membranes with 1 μm-wide pores in the presence of an apicobasal EGF gradient. After 12 hours, GFP-SEPT9_1 localized in the shaft and the base of shorter invadopodia, forming a ring-like structure (Figure 1D-E). Three-dimensional rendering showed that GFP-SEPT9_i1 decorated the basal sides of a subset of invadopodia, forming a barrel-like structure at the proximal end of invadopodia (Figure 1E). After 24 h of EGF treatment, GFP-SEPT9_i1 became more enriched in the distal segments of elongated mature invadopodia (Figure 1F-H, arrows). In ∼40% of these invadopodia, GFP-SEPT9_i1 localized exclusively in their distal ends and tips, which indicated a shift in the localization of GFP-SEPT9_i1 during invadopodia maturation. Collectively, these data suggest that SEPT9_i1 is an integral component of invadopodia, and may function in invadopodia formation and maturation.

### SEPT9_i1 is required for invadopodia formation and ECM degradation

To test whether SEPT9_i1 is specifically required for the formation of functional invadopodia, we generated lentiviral vectors with shRNAs that target unique sequences of the SEPT9 isoforms SEPT9_i1 (shRNA-1) and SEPT9_i2 (shRNA-2), and a sequence that is shared among all SEPT9 isoforms (shRNA-3). We chose to target SEPT9_i2, because it is an isoform that inhibited the migration of breast cancer MCF-7 cells in a 3D transwell assay, and thus may functionally differ from SEPT9_i1 ^33^. SEPT9_i2 has a distinct N-terminal sequence, which lacks the first 25 amino acids of SEPT9_i1 and is encoded by an alternative exon that is hypermethylated in breast and colon cancers ^33, 44, 45^. Unlike SEPT9_i1, which associates strongly with actin filaments ^34, 41, 46^, SEPT9_i2 colocalizes weakly with actin, and SEPT9_i2 over-expression reduces septin association with actin filaments ^33^.

Staining MDA-MB-231 cells with isoform-specific antibodies against SEPT9_i1 and SEPT9_i2 showed that their expression was knocked-down by shRNA-1 and shRNA-2, respectively (Figure 2A-D). Similar to shRNA-1, shRNA-3 also reduced SEPT9_i1 expression (Figure 2A, C). SEPT9_i1-depleted cells were devoid of ventral and perinuclear septin filaments, and they had background-levels of fluorescence intensity with no discernable structures (Figure 2A-B). As previously reported, SEPT9_i2 had fewer and shorter filaments than SEPT9_i1 ^33^, which dissipated in cells treated with shRNA-2 (Figure 2B). Consistent with previous observations ^41, 42, 47^, SEPT9 knock-down disrupted the filamentous organization of SEPT2 and SEPT7 on both the ventral and dorsal sides of the nucleus (Figure S2A-D). SEPT2 and SEPT7 were nearly absent from the ventral surface of the nucleus, and filaments were reduced to puncta in the mediodorsal regions of the cytoplasm around the nucleus (Figure S2A-D).

**Figure 2.**
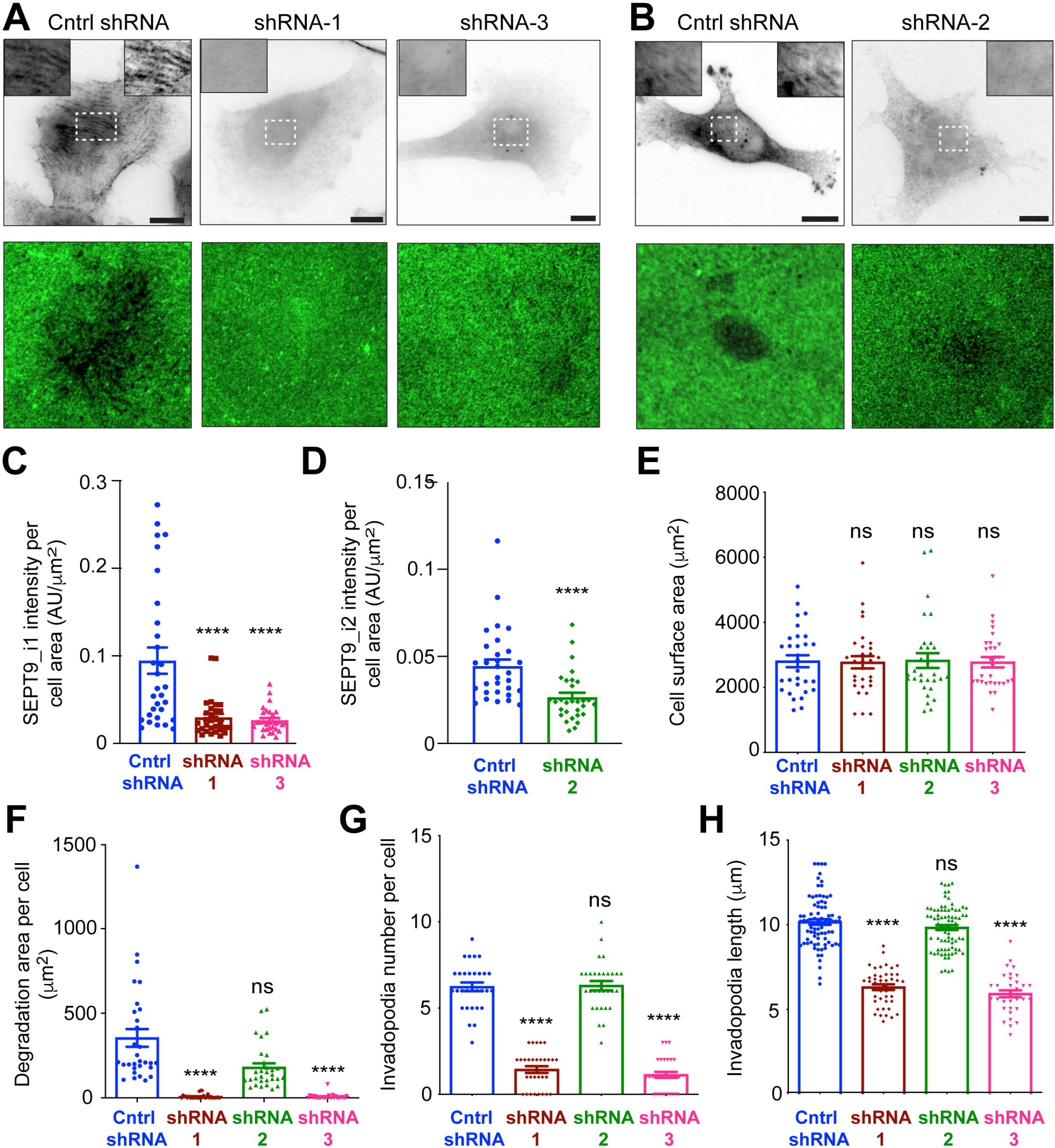
SEPT9_i1 depletion abrogates the formation of ECM degrading invadopodia. (A-B) MDA-MB-231 cells were transduced with lentiviruses carrying control shRNAs and shRNAs targeting SEPT9_i1 (shRNA1), SEPT9_i2 (shRNA2) or all SEPT9 isoforms (shRNA3), and were plated on gelatin-AlexaFluor594 matrix for 5 h in the presence of EGF (10 nM for 4h). Images show cells stained with isoform-specific antibodies against SEPT9_i1 (A; inverted monochrome) or SEPT9_i2 (B; inverted monochrome) and the underlying gelatin matrix (green). Insets show in higher magnification areas outlined with dashed rectangles. Right corner insets show SEPT9 filaments in higher contrast. Scale bars, 10 μm. (C) Quantification of SEPT9_i1 fluorescence (mean ± SEM) in MDA-MB-231 cells, which were transduced for 60 h with lentivirus carrying control shRNAs (n = 30) or shRNAs targeting SEPT9_i1 (shRNA 1; n = 30) or all SEPT9 isoforms (shRNA 3; n = 30). (D) Quantification of SEPT9_i2 fluorescence (mean ± SEM) in MDA-MB-231 cells, which were transduced with lentivirus carrying control shRNAs (n = 29) or SEPT9_i2 shRNA (shRNA2; n = 29). (E) Cell surface area (mean ± SEM) of MDA-MB-231 cells, which were transduced for 60 h with lentivirus carrying control shRNAs (n = 30) or shRNAs targeting SEPT9_i1 (shRNA1; n = 30), SEPT9_i2 (shRNA2; n = 30) or all SEPT9 isoforms (shRNA3; n = 30). (F) Quantification of the surface area of gelatin-AlexaFluor594 that was degraded per MDA-MB-231 cell (mean ± SEM; n = 30). Prior to plating on gelatin for 4 h in EGF, MDA-MB-231 cells were treated with shRNAs for 48 h. (G-H) Quantification of the mean (± SEM) number of invadopodia per cell (n = 30; G) and mean length of invadopodia (n = 34 – 83; H) in MDA-MB-231 cells, which were plated for 24 h on 1 μm-pore size transwell membranes with an EGF gradient after infection and selection for shRNA expression. Data were analyzed with either a non-parametric one-way ANOVA Kruskal-Wallis test with Dunn’s post-hoc analysis for pairwise comparisons (C, E-G) or a one-way Welch ANOVA test with Dunnett T3 post-hoc test for pairwise analysis with control shRNAs (H). Statistical analysis in (D) was performed with the Mann-Whitney U-test. ns, not significant; ****, p < 0.0001

In 2D assays of fluorescent gelatin degradation, we treated MDA-MB-231 cells for 4 h with EGF after ∼60 h of SEPT9 depletion. Knock-down of SEPT9_i1 by shRNA-1 and shRNA-3, which targets all SEPT9 isoforms, reduced drastically the degradation of gelatin without affecting cell surface areas (Figure 2E-F). In contrast, depletion of SEPT9_i2 did not impact gelatin degradation (Figure 2E-F). Consistent with these results, invadopodia formation was also markedly reduced in 3D chemoinvasion assays (Figure 2G-H). SEPT9_i1 knock-down by shRNA-1 and shRNA-3 diminished the mean number of invadopodia per cell (Figure 2G). In addition, the mean length of invadopodia diminished (Figure 2H), indicating that SEPT9_i1 is critical for both the formation and the elongation of invadopodia. SEPT9_i2 knock-down, however, did not alter the number or length of invadopodia (Figure 2G-H). To further test whether the SEPT9_i1 isoform was sufficient in inducing invadopodia and to demonstrate that the effects of septin depletion were specific, we sought to rescue invadopodia reduction with an shRNA-resistant SEPT9_i1^res^-GFP (SEPT9_i1^res^-GFP). In the 3D chemoinvasion assay, SEPT9_i1^res^-GFP restored the number of invadopodia, which was markedly reduced with shRNA that targeted all SEPT9 isoforms (Figure S2E-F). Taken together, these data show that the oncogenic SEPT9 isoform 1 promotes invadopodia formation and invadopodia-mediated ECM degradation.

### SEPT9_i1 depletion disrupts the clustering and localization of the invadopodia precursor components TKS5 and cortactin

Because SEPT9_i1 depletion impairs invadopodia formation, we examined whether SEPT9_i1 is functionally involved in the assembly of the invadopodia precursor complexes which begins with the recruitment and clustering of cortactin and TKS5 on domains of the ventral plasma membrane ^9, 13, 48, 49^. We first stained for the scaffold protein TKS5 in MDA-MB-231 cells, which were cultured on gelatin in the presence of EGF. In ventral sections of control cells, TKS5 had a punctate appearance, forming clusters of variable size that localize to areas of the plasma membrane underneath and around the nucleus (Figure 3A). In cells treated with shRNAs that target SEPT9_i1 (shRNA-1) or all SEPT9 isoforms (shRNA-3), there was a striking reduction in the number and size of TKS5 puncta (Figure 3A-C). TKS5 had a more granular appearance with smaller puncta outlining the nuclear rim, which was suggestive of TKS5 dispersion and mislocalization along the nuclear envelope (Figure 3A). The lower intensity and smaller size of these puncta resembled background staining, and they were below the threshold range of TKS5 cluster segmentation (outlined clusters in Figure 3A insets). In contrast, knock-down of SEPT9_i2 with shRNA-2 did not disrupt TKS5 clusters and TKS5 localization was similar to control cells (Figure 3A-C).

**Figure 3.**
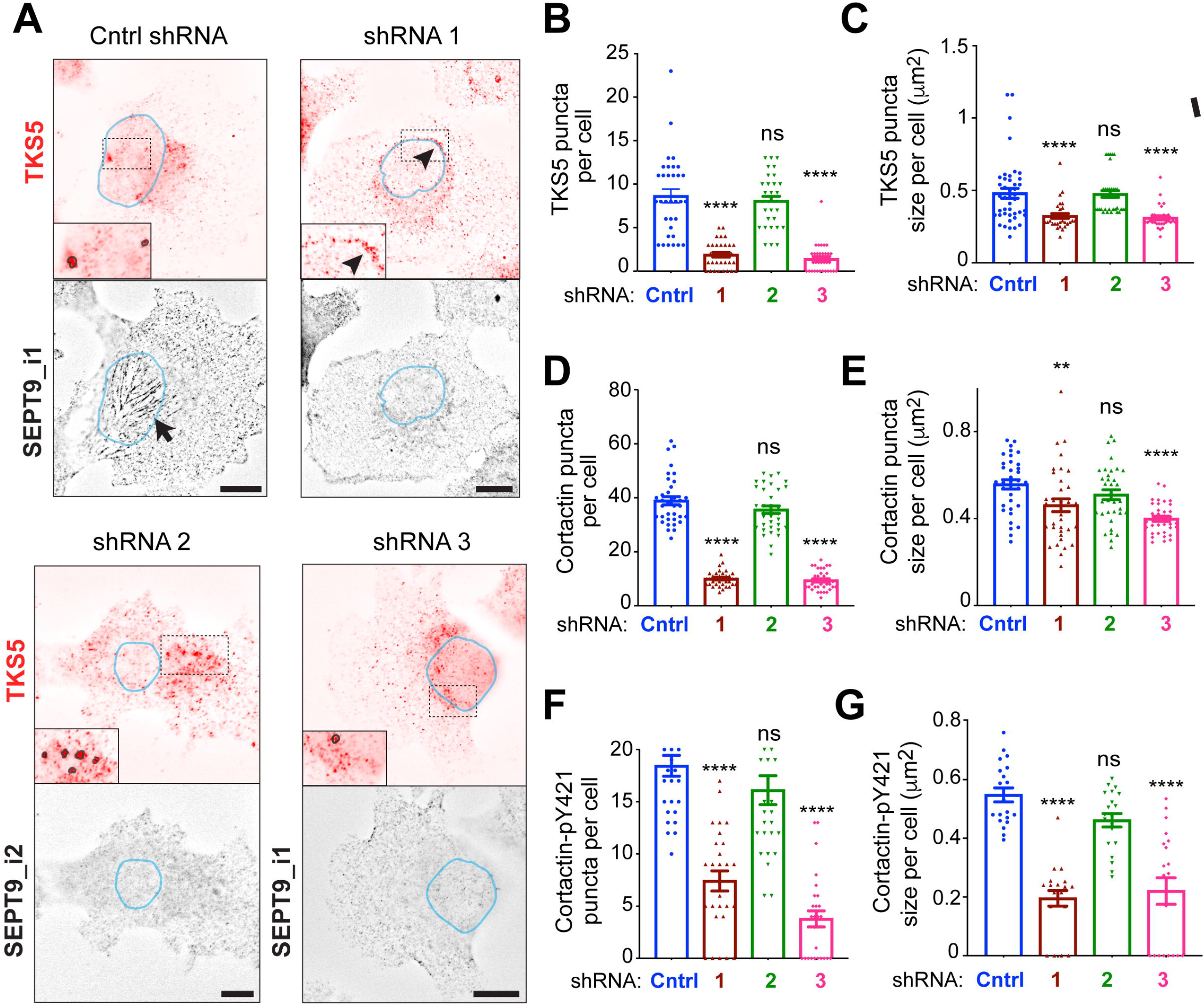
SEPT9_i1 depletion reduces the number and size of the ventral clusters of TKS5, cortactin and cortactin-pY421. (A) Images of the ventral surface of MDA-MB-231 cells, which were treated with shRNAs for 48 h, and stained for TKS5, SEPT9_i1 and SEPT9_i2. Insets show the areas outlined with dashed lines in higher magnification, and blue lines outline the nuclear regions. In insets, encircled puncta outline the TKS5 clusters which met the segmentation thresholding range applied for their identification and quantification. Arrowhead points to diminished TKS5 granules outlining the nuclear rim. Arrow points to ventral SEPT9_i1 filaments, which are absent from cells treated with SEPT9 shRNAs. All images were processed with constrained iterative deconvolution to remove out of focus light. Scale bars, 10 μm. (B-C) Bar graphs show the mean (± SEM) number (B) and size (C) of ventral TKS5 puncta in MDA-MB-231 cells (n = 30-36) treated with control shRNAs and shRNAs against SEPT9_i1 (shRNA1), SEPT9_i2 (shRNA2) and all SEPT9 isoforms (shRNA3). (D-E) Bar graphs show the mean (± SEM) number (D) and size (E) of ventral cortactin puncta in MDA-MB-231 cells (n = 35) treated with control shRNAs and shRNAs against SEPT9_i1 (shRNA1), SEPT9_i2 (shRNA2) and all SEPT9 isoforms (shRNA3). (F-G) Bar graphs show the mean (± SEM) number (F) and size (G) of ventral cortactin-pY421 in MDA-MB-231 cells (n = 27) treated with control shRNAs and shRNAs against SEP9_i1 (shRNA1), SEPT9_i2 (shRNA2) and all SEPT9 isoforms (shRNA3). Data were statistically analyzed with the Kruskal-Wallis one-way ANOVA test and the p-values were derived with Dunn’s post-hoc analysis for pairwise comparisons with control shRNA. ns, not significant; **, p < 0.01; ****, p < 0.0001

Next, we examined whether SEPT9_i1 depletion had a similar effect on the localization of cortactin and phosphorylated cortactin-pTyr421, which promote actin polymerization and the maturation of nascent invadopodia. In control MDA-MB-231 cells, cortactin and cortactin-pTyr421 puncta were more numerous than TKS5, but their mean size was similar to TKS5 puncta (Figure 3D-G). Depletion of SEPT9_i1 and all SEPT9 isoforms markedly reduced the number of cortactin and cortactin-pTyr421 puncta per cell, while knock-down of SEPT9_i2 had no impact (Figure 3D-G). These data are consistent with the reduction of TKS5, which is recruited to cortactin/actin-containing invadopodia precursors ^13^. Therefore, the oncogenic SEPT9_i1 isoform is required for the assembly of invadopodia precursor complexes into plasma membrane clusters, and their maturation into TKS5-positive invadopodia.

### SEPT9_i1 localizes to the basolateral surfaces of the nuclear envelope and juxtanuclear invadopodia

Previous work showed that the apical ends of invadopodia physically contact the nuclear envelope, which in turn may counteract the forces that resist invadopodia elongation and protrusion into the matrix ^23^. We reasoned that defective assembly and maturation of invadopodia might be due to due to lack of mechanical support by the nuclear envelope. To investigate this possibility, we examined first whether SEPT9_i1 localizes to the nuclear envelope and/or any invadopodia that are juxtaposed to the nucleus.

Septin filaments have been reported to localize at subnuclear and perinuclear regions of the plasma membrane and the cytoplasm in a variety of cell types ^50, 51^. However, it is unknown whether septins colocalize with the nuclear envelope. Using high- and super-resolution confocal microscopy, we examined the localization of endogenous SEPT9_i1 with respect to the nuclear envelope of MDA-MB-231 cells, which was labeled with antibodies to lamin A/C and nucleoporins. In confocal super-resolution sections, which were serially acquired in the ventrodorsal direction of lamin A/C-stained nuclei, SEPT9_i1 filaments transversed the ventral sections of the nucleus and gradually disappeared in sections that were over ∼200 nm from the nuclear surface (Figure 4A). Three-dimensional rendering of super-resolution confocal images of nucleoporin and SEPT9_i1 showed that SEPT9_i1 filaments flank nucleoporin-labeled puncta (Figure 4B-C), indicating that SEPT9_i1 is closely apposed to the nuclear envelope. In mediobasal confocal sections, SEPT9_i1 filaments also co-aligned with segments of the nuclear rim, which was labeled with lamin A/C (Figure 4D).

**Figure 4.**
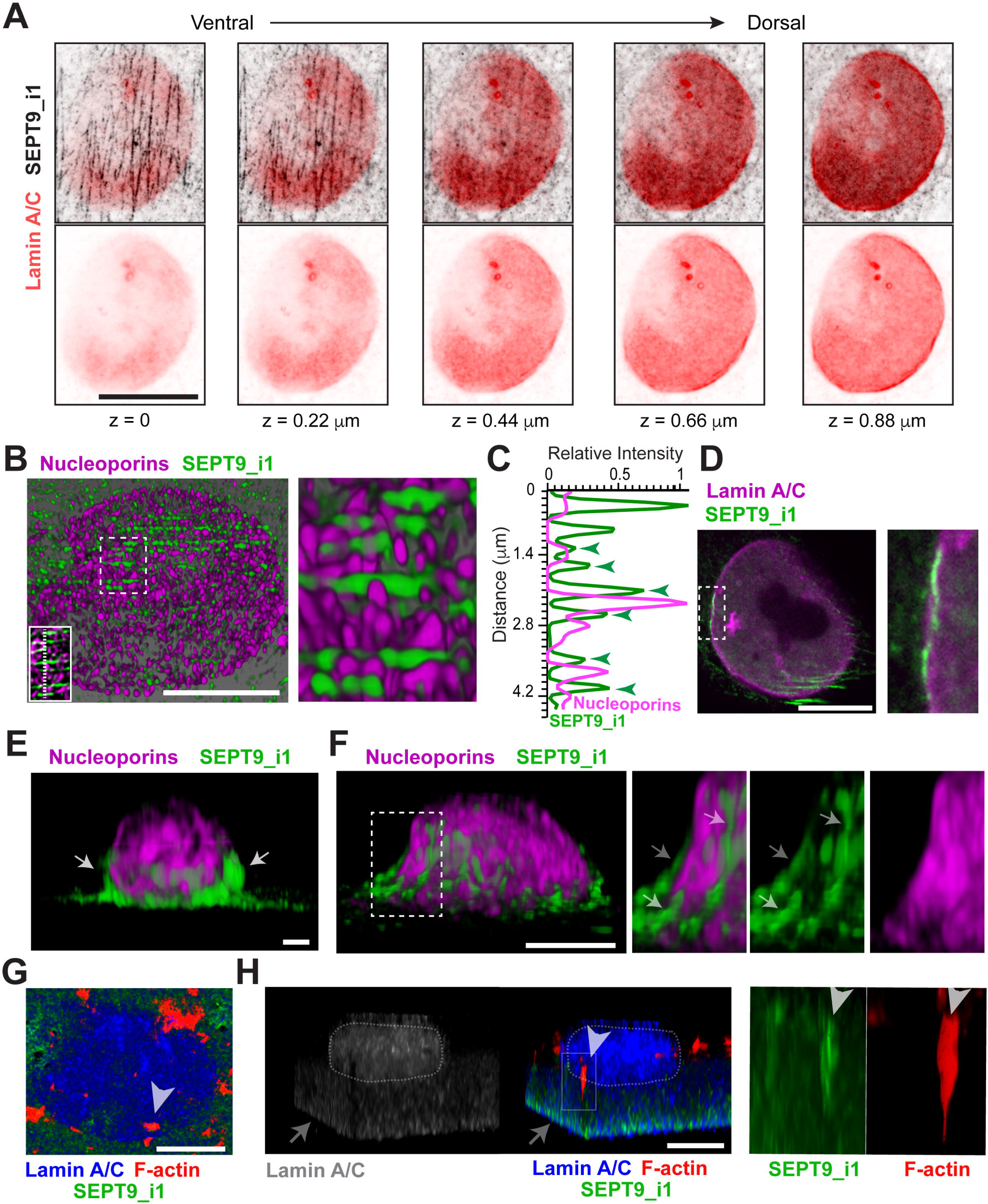
SEPT9_i1 localizes to the basolateral surfaces of the nuclear envelope, and colocalizes with juxtanuclear invadopodia. (A) Super-resolution confocal microscopy sections of an MDA-MB-231 cell, which was stained for endogenous SEPT9_i1 (inverted monochrome; black) and lamin A/C (burgundy) after plating on gelatin for 5 h and treated with EGF for 4 h. Image series shows consecutive images, which were taken in the ventral-dorsal direction at a step size of 0.22 μm, demonstrating that SEPT9_i1 localizes to the ventral surface of the nuclear envelope. Scale bar, 5 μm. (B) Image shows a 3D view of super-resolution confocal microscopy sections, which were taken from the ventral side of the nuclear envelope of an MDA-MB-231 cell that was stained for endogenous nucleoporins (mAb 414) and SEPT9_i1. Outlined area (dashed rectangle) is shown in higher magnification. Inset shows confocal microscopy section of the same region; dashed line indicates the region of the fluorescence intensity plot profile in panel C. Scale bar, 5 μm. (C) Fluorescence intensity plot profile of the fluorescence of SEPT9_i1 (green) and nucleoporins (magenta) across the dashed line shown in the inset of panel B. Arrowheads point to fluorescence intensity peaks of SEPT9_i1 filaments that flank nucleoporins. (D) Spinning disk microscopy image of the nuclear area of an MDA-MB-231 cell, which was plated on gelatin and stained for endogenous lamin A/C and SEPT9_i1. In the selected area, which is shown in higher magnification, a SEPT9_i1 filament is closely juxtaposed to the nuclear rim. Scale bar, 5 μm. (E-F) Images show 3D views of super-resolution confocal microscopy sections of MDA-MB-231 cells which were plated on collagen and treated with EGF for 4 h. Cells were stained for endogenous nucleoporins (mAb 414) and SEPT9_i1. Arrows point to SEPT9_i1 filaments along the mediolateral sides of the nucleus (E) and SEPT9_i1 filaments with concave curvature (F) stretching along the nuclear surface. Scale bars, 1 μm (E) and 5 μm (F). (G-H) Top-down (G) and side (H) views of 3D rendered super-resolution confocal microscopy images of lamin A/C, SEPT9_i1 and actin (phalloidin) in an MDA-MB-231 cell, which was plated on a matrigel- and laminin-coated Transwell membrane with an EGF gradient. The lamin-stained mass of the nucleus is outlined with a dashed line. Arrowhead points to an invadopodium that extends from the lateral surface of the nucleus toward the plasma membrane. Arrow points to the Transwell membrane, which is visible due to non-specific background stain. Dashed rectangle encloses and invadopodium shown in higher magnification on the right. Scale bars, 5 μm.

To enhance our view of SEPT9_i1 organization around the 3D mass of the nucleus and its envelope, we examined the localization of SEPT9_i1 in 3D images of cells with nuclei which had a more spherical than oblate shapes. Sagittal views of 3D-rendered super-resolution confocal sections revealed SEPT9_i1 filaments, which were oriented vertically along the mediobasal curvature of the nuclear envelope (Figure 4E). Several SEPT9_i1 filaments had a concave-shaped curvature and stretched along the downward slope of the nucleoporin-labeled nuclear envelope (Figure 4F, arrows). The orientation and negative bent of these SEPT9_i1 filaments were similar to the organization of septins on sinusoidal substrates, on which septins form filaments that bend along concave curvatures ^52^.

The apicobasal orientation of SEPT9_i1 filaments along the lateral curvature of the nuclear envelope prompted us to examine how SEPT9_i1 localized with respect to invadopodia that contact the nuclear envelope. We performed 3D confocal super-resolution microscopy of MDA-MB-231 cells cultured on Matrigel- and laminin-coated transwell membranes with an EGF gradient. Three-dimensional rendering of these cells showed the position of the lamin A/C-stained nucleus with respect to the porous membrane, which is visible due to background staining (Figure 4G-H). In side-views of the nucleus, we observed actin-rich needle-shaped structures with a wider apical end and a narrowing thinner basal tip (Figure 4H), which resembled the nucleus-bound invadopodia described by correlative fluorescence microscopy and focus ion beam-scanning electron microscopy ^23^. SEPT9_i1 was enriched in the segment that overlapped with the nucleus, indicating that SEPT9_i1 is present in invadopodia that are juxtaposed to the nucleus (Figure 4G-H, arrowheads). Collectively, these imaging data indicate that SEPT9_i1 localizes to the ventral and mediobasal surfaces of the nuclear envelope, and juxtanuclear invadopodia.

### SEPT9_i1 knock-down causes nuclear envelope deformities, altering nuclear shape and volume

We next asked whether SEPT9_i1 influences nuclear morphology by knocking down expression of SEPT9_i1 or all SEPT9 isoforms. Confocal 3D imaging of lamin A/C and nucleoporin revealed that the nuclei of MDA-MB-231 cells were strikingly altered (Figure 5A-C). Nuclear shapes were compressed and irregular, and the nuclear envelope was herniated containing multiple folds and indentations (Figure 5A). In contrast, cells treated with control shRNA contained oval-shaped nuclei with a smooth nuclear envelope (Figure 5A). In 3D-rendered images, these nuclei were flatter than those of SEPT9_i1-depleted cells, which were taller and crumpled with multiple creases (Figure 5B, arrows). Throughout the nucleus, we observed grooves extending inward from indentations of the nuclear envelope (Figure 5B). These nuclear grooves were rich in lamin A/C and nucleoporins (Figure 5C-F), which are indicative of nuclear envelope invaginations and folds. In addition to the nuclear deformities, perinuclear actin filaments were diminished (Figure S3A-D), which was consistent with previous loss of actin stress fibers in cells depleted of SEPT9 ^41, 42, 47^. Notably, there was an amplification of actin patches in the mediodorsal sections of the nucleus, several of which localized at regions of the distorted nuclear rim (arrows in Figure S3B, D).

**Figure 5.**
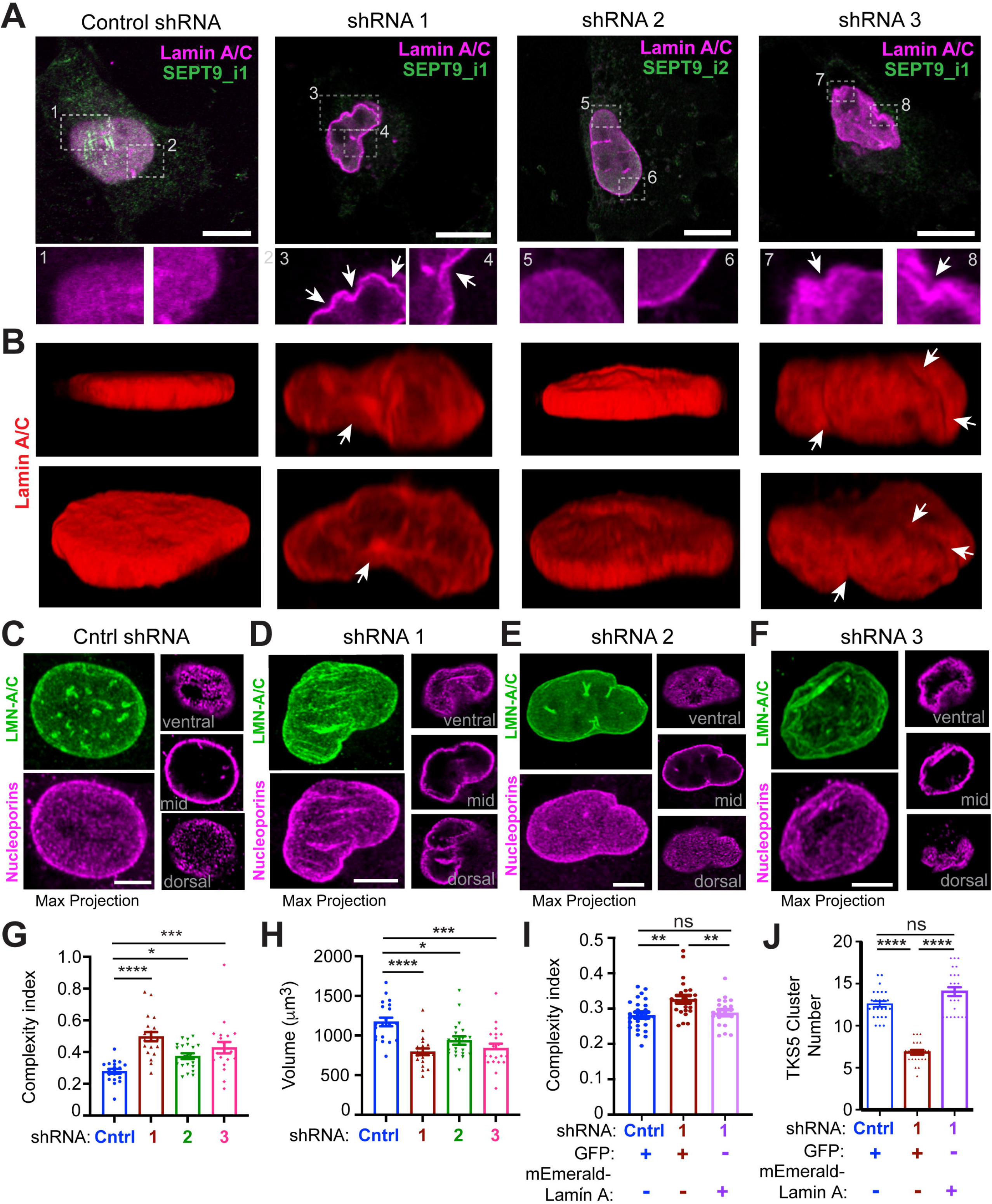
SEPT9_i1 depletion causes nuclear envelope folds and grooves, altering nuclear shape and volume. (A) Confocal microscopy images of MDA-MB-231 cells, which were stained for endogenous lamin A/C (magenta) and SEPT9_i1 or SEPT9_i2 (green) after treatment with shRNAs and plating on gelatin. Areas outlined in numbered dashed rectangles are shown in higher magnification. Arrows point to folds and herniations of the nuclear envelope. Scale bars, 5 μm. (B) Side-views of 3D rendered confocal images of the lamin A/C-stained nuclei from the cells and conditions, which are shown above in (A). Arrows point to folds and herniations of the nuclear envelope. (C-F) MDA-MB-231 were plated on gelatin and treated with control shRNAs (C) and shRNAs targeting SEPT9_i1 (D; shRNA1), SEPT9_i2 (E; shRNA 2) and all SEPT9 isoforms (F; shRNA 3). Cells were stained for endogenous lamin A/C (green) and nucleoporins (magenta). Images show maximum projections of super-resolution confocal microscopy sections (left), and single optical slices from the ventral, middle and dorsal sections of each z-series stack (right). Nuclear envelope folds and herniations are evident by nucleoporin stain in cells treated with shRNA 1 (D) and shRNA 3 (F). Arrows point to nuclear grooves and folds. Scale bars, 5 μm. (G-H) Quantification of the mean (± SEM) nuclear complexity index (ratio of nuclear perimeter to surface area; G) and volume (H) in MDA-MB-231 cells treated for 48 h with control shRNAs (n = 20) and shRNAs targeting SEPT9_i1 (shRNA1; n = 20), SEPT9_i2 (shRNA 2; n = 20) and all SEPT9 isoforms (shRNA 3; n = 20). Data in (G) were analyzed with a non-parametric Kruskal-Wallis one-way ANOVA with Dunn’s post-hoc analysis for pair-wise comparisons, and data in (H) with a Welch ANOVA test with Dunnett T3 post-hoc analysis for pair-wise comparisons with control shRNA. (I-J) Quantification of the mean (± SEM) nuclear complexity index (ratio of nuclear perimeter to surface area; I, n = 21-26) and ventral TKS5 puncta per cell (J, n = 23-25) in MDA-MB-231 cells, which were transduced with viruses carrying shRNAs for 48 h, and subsequently transfected with GFP or mEmerald-lamin A for 48 h. Statistical analysis of pair-wise comparisons was performed with a Mann-Whitney U-test or Welch’s t-test. ns, not significant; *, p < 0.05; **, p < 0.01; ***, p < 0.001; ****, p < 0.0001

Quantifications of nuclear morphology were consistent with the images of deformed nuclei. The nuclear complexity factor, which is the ratio of the nuclear perimeter to surface area, increased in cells depleted of SEPT9_i1 and all SEPT9 isoforms (Figure 5G). Nuclear volumes also decreased (Figure 5H). Changes in these parameters reflected the wrinkling of the nuclear envelope and crumpling of the nuclear mass, which resulted in smaller nuclei with zigzagged perimeters. In contrast, the nuclei of SEPT9_i2-depleted cells were largely oval and flat but contained a few creases and wrinkles (Figure 5B). This phenotype was quantitatively and statistically weaker than SEPT9 depletion with shRNAs 1 and 3 (Figure 5G-H). These results reveal that in SEPT9_i1-depleted cells, defective invadopodia assembly is accompanied with nuclear deformation, while SEPT9_i2 depletion has a mild effect on nuclear morphology with no impact on invadopodia.

The nuclear deformations of SEPT9_i1 depletion phenocopy those of laminopathies, in which nuclear lamins are mutated ^53, 54^. Because lamins promote the mechanical stability and stiffness of the nucleus ^55–57^, we reasoned that SEPT9_i1 may function similar to nuclear lamins in promoting nuclear stability. To test this hypothesis, we asked first whether SEPT9_i1 depletion affects lamin A expression, second if over-expression of lamin A can counteract the nuclear deformities of SEPT9_i1 depletion, and third whether the reverse is possible. Quantification of endogenous lamin A levels across the z-axis of the nucleus showed no effect by SEPT9 shRNAs (Figure S3E). In agreement, previous work found that SEPT9_i1 over-expression had no impact on lamin A expression in MCF7 breast cancer cells ^36^. We then expressed mEmerald-lamin A in MDA-MB-231 cells, which were treated with shRNA1, and stained for nucleoporins to compare nuclear morphology to controls - cells that were transfected with GFP after treatments with either control shRNA or shRNA1. To avoid any artifacts from over-expression of mEmerald-lamin A, which can aggregate in the nucleoplasm, we only imaged cells with moderate levels of mEmerald-lamin A expression. Lamin A rescued the nuclear phenotype of SEPT9_i1 knock-down, decreasing the nuclear complexity factor, which was not statistically different from that of cells treated with control shRNAs (Figure 5I). Conversely, expression of GFP-tagged SEPT9_i1, which rescued the nuclear complexity factor in SEPT9_i1-depleted cells, ameliorated the nuclear defects of lamin A knock-down (Figure S4). Thus, SEPT9_i1 appears to function redundantly with lamin A in promoting the mechanical stability of the nucleus.

Given the strong correlation between nuclear deformation and loss of invadopodia assembly, we tested if mEmerald-lamin A expression affected invadopodia formation on 2D matrices of gelatin. Indeed, mEmerald-lamin A restored the number of TKS5 clusters to control levels (Figure 5J). In the absence of a confined mode of migration, rescue of TKS5 clustering suggests that invadopodia assembly is coupled to the nucleus independently of an adaptive response to a dense ECM. Moreover, the oncogenic SEPT9_i1 appears to regulate a crosstalk between the mechanical properties of the nucleus and the machinery of invadopodia formation.

### SEPT9_i1 is required for EGF-dependent enhancement of juxtanuclear invadopodia

Invadopodia formation has been reported to occur preferentially in plasma membrane regions that overlap with the nucleus ^19, 23^, but how this spatial bias is generated is little understood. We sought to examine whether SEPT9_i1 contributes to the preferential formation of juxtanuclear invadopodia. We first analyzed the subcellular distribution of invadopodia and SEPT9_i1 before and after treatment with EGF, which activates invadopodia formation. We analyzed the subcellular distribution of ventral clusters that contained actin, TKS5 and cortactin-pY421, which is a marker of invadopodium maturation ^9, 58, 59^. In MDA-MB-231 cells, which were plated on non-fluorescent gelatin, we quantified the percentage of ventral actin/TKS5/cortactin-pY421-containing clusters that overlapped with the nuclear footprint and a 3 μm-wide zone around the nucleus. EGF enhanced invadopodia formation within the nuclear footprint, including invadopodia that were positioned on the nuclear rim (Figure 6A, arrows), increasing the percentage of juxtanuclear invadopodia by two-fold (Figure 6B). In parallel with this increase in maturing invadopodia, EGF also amplified the overlap of ventral SEPT9_i1 filaments with the nucleus (Figure 6C-D). The percentage of the nuclear area that overlapped with SEPT9_i1 underwent a near two-fold increase (Figure 6B). Thus, EGF not only induces the formation of invadopodia, but also biases their formation in juxtanuclear regions with concomitant amplification of SEPT9_i1 in proximalmost areas of the nuclear envelope.

**Figure 6.**
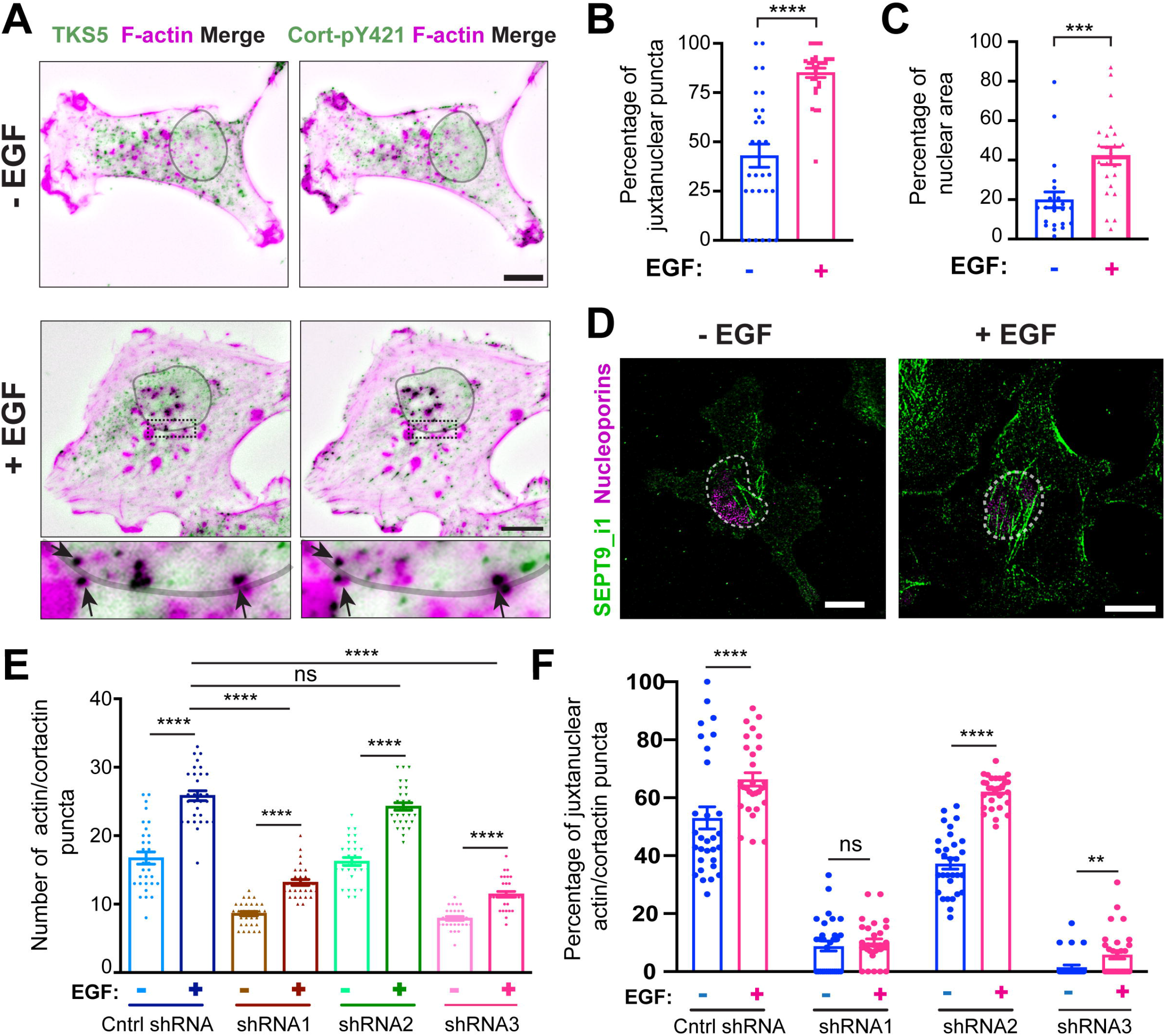
SEPT9_i1 is required for EGF-induced amplification of juxtanuclear invadopodia. (A-B) Images (A) show the localization of ventral TKS5 puncta with respect to the nuclear area (DAPI) in MDA-MB-231 cells, which were plated on gelatin with or without EGF treatment. Bar graph (B) shows the number of TKS5 puncta, which overlap with the DAPI-stained nuclear footprint area and a 3 μm-wide zone around the nucleus, as percentage of total TKS5 puncta in MDA-MB-231 cells (n = 32). Arrows point to TKS5/actin/cortactin-pY421 clusters that localize at the nuclear rim. Data were statistically analyzed with an unpaired t-test. Scale bars, 10 μm. (C-D) Bar graph (C) shows the percentage of the nuclear footprint area that SEPT9_i1 occupies in MDA-MB-231 cells (n = 22-23) after a 4 h incubation with or without EGF. Data were statistically analyzed with the Mann-Whitney U-test. Super-resolution confocal microscopy images (D) show the localization of ventral SEPT9_i1 with respect to the nuclear area (nucleoporins) in MDA-MB-231 cells with and without EGF treatment. (E) Bar graphs show quantifications of the number of ventral cortactin-positive actin puncta in MDA-MB-231 cells (n = 30), which were treated with shRNAs for 48 h and incubated with or without EGF for 4 h. Statistics for pairwise comparisons were derived with the Welch’s t-test (control shRNA), Mann-Whitney U-test (shRNA1 and shRNA3), and student’s t-test (shRNA2). (F) Bar graphs show number of ventral cortactin-positive actin puncta which overlap with the nuclear footprint area and a 3 μm-wide zone around the nucleus as percentage of the total ventral cortactin-positive actin puncta. Quantifications were performed in MDA-MB-231 cells (n = 30), which were treated with shRNAs for 48 h and incubated with or without EGF for 4 h. Statistics for pairwise comparisons were derived with the Welch’s t-test (control shRNA, shRNA2) and Mann-Whitney U-test (shRNA1 and shRNA3). ns, not significant; **, p <0.01; ***, p < 0.001; ****, p < 0.0001

To test whether SEPT9_i1 is required for the EGF-induced formation of juxtanuclear invadopodia, we analyzed the formation and distribution of ventral invadopodia precursors with or without EGF treatment in control and SEPT9 depletion background. Consistent with the data above (Figure 3), depletion of SEPT9_i1 with shRNAs 1 and 3 diminished the number of invadopodia precursor clusters at steady state and upon EGF treatment (Figure 6E and Figure S5A-D). By contrast, invadopodia precursor numbers were not statistically different between control and SEPT9_i2 knock-down cells (Figure 6E). In SEPT9_i1-depleted cells, EGF enhanced the number of invadopodia precursors in spite of being lower than matched controls (Figure 6E). Thus, SEPT9_i1 depletion does not impede the upregulation of invadopodia precursors by EGF. However, analysis of the percentage of ventral precursors that overlap with the nuclear footprint revealed a drastic elimination of juxtanuclear precursors, which were reduced from 50%-60% to 4-5% of total precursors (Figure 6F and S5A-D). This dearth of juxtanuclear invadopodia persisted in the presence of EGF. In MDA-MB-231 cells treated with shRNA1, EGF did not increase juxtanuclear invadopodia, and in cells treated with shRNA3, EGF did not raise the percentage of juxtanuclear invadopodia above 4% (Figure 6F). To the contrary, EGF enhanced the percentage of juxtanuclear invadopodia precursors in control and SEPT9_i2-depleted cells (Figure 6F). Notably, expression of mEmerald-lamin A in SEPT9_i1-depleted MDA-MB-231 cells enhanced the percentage of ventral juxtanuclear TKS5 puncta (Figure S5E), which indicated that the juxtanuclear positioning of invadopodia in response to EGF is dependent on the mechanical strength of the nucleus.

Given the use of the MDA-MB-231 cell model throughout this study, we asked whether SEPT9_i1 functions similarly in the 786-0 renal carcinoma cells. Endogenous SEPT9_i1 localized as filaments across the ventral surface of the nucleus (Figure S6A). SEPT9 knock-down resulted in loss of ventral SEPT9_i1 filaments (Figure S6A), and decreased the degradation of fluorescent gelatin, a phenotype that was rescued by SEPT9_i1^res^-GFP (Figure S6B). In 786-0 cells, SEPT9 knock-down reduced the total number TKS5 puncta and the percentage of ventral juxtanuclear TKS5 puncta, which increased upon expression of SEPT9_i1^res^-GFP (Figure S6D-F). Consistent with the role of SEPT9_i1 in the nuclear stability of MDA-MB-231 cells, SEPT9 knock-down increased the number of nuclear grooves in 786-0 cells, and complementation of SEPT9 depletion with SEPT9_i1^res^-GFP suppressed the formation of nuclear grooves (Figure S6G-H). Taken together, these results indicate that SEPT9_i1 promotes nuclear envelope stability and biases invadopodia formation at juxtanuclear regions of the plasma membrane in cancer cells of different tissue origin.

## DISCUSSION

Cancer cell migration and invasion are largely guided by the biophysical properties of the ECM and the nucleus, and their mechanical crosstalk ^17, 18, 60^. Invadopodia enable nuclear passage by degrading the ECM, and constitute an invasive strategy that is favored by nuclei of low deformability and ECMs of high density ^19–21^. Although the nucleus is increasingly viewed as a mechanosensory organelle that determines migratory strategies in response to outside-in forces, it is unknown if and how the nucleus crosstalks with invadopodia. Previous work suggested a physical link between the nuclear envelope with the actin core of invadopodia, which may counterbalance the forces exerted on the distal end of invadopodia by the ECM ^23^. New evidence, however, shows that invadopodia exert outward forces that push and remodel ECM fibers through actin polymerization ^24^. Therefore, it is unclear whether invadopodia encounter substantial resistance forces, which are transmitted and counterbalanced by the nucleus.

In this study, our results demonstrate that invadopodia formation is spatially biased, taking place preferentially at plasma membranes sites which are juxtaposed to the nuclear envelope. Importantly, our data indicate that nuclear stability is critical for invadopodia formation, and for the spatiofunctional coupling of invadopodia to the nucleus. We identified the oncogenic isoform 1 of SEPT9 as an essential and novel component of invadopodia, which links functionally the machinery of invadopodia formation to the nuclear envelope. Collectively, our data have revealed a hitherto unknown and isoform-specific function for SEPT9_i1 in nuclear morphology and invadopodia formation.

SEPT9_i1 over-expression has been linked to a variety of oncogenic processes including cell proliferation ^31^, genomic instability ^61^, hypoxia-induced angiogenesis ^32, 62^, and cell migration ^27, 33–35^. In the context of cell migration and invasion, SEPT9_i1 has been mechanistically linked to focal adhesion maturation and stability ^34, 35^. In breast cancer cells, SEPT9 isoform 3 enhances the secretion of MMPs at focal adhesions ^36^, but no SEPT9 isoforms or other septins have been previously implicated in the degradation of ECM by invadopodia. In endothelial cells, however, SEPT2 promotes the maturation of podosomes by forming a putative diffusion barrier around membrane-bound TKS5- and actin-containing complexes ^40^.

In contrast to the function of SEPT2 in podosomes ^40^, SEPT9_i1 influences the initial clustering of invadopodia precursor components such as cortactin and TKS5, and spatially biases their localization to juxtanuclear regions of the plasma membrane. Although we cannot rule out the possibility of SEPT9_i1 playing a role in the plasma membrane, SEPT9_i1 appears to impact invadopodia formation by bolstering the mechanical stability of the nuclear envelope, as evidenced by rescue of the reduction of TKS5 puncta by lamin A in SEPT9_i1-depleted cells. We posit that in early stages of invadopodia formation, SEPT9_i1 may promote invadopodia assembly at membrane contact sites between the nuclear envelope and cell cortex. In HeLa cells, septins localize at junctions of the plasma membrane with the ER, which is contiguous with the nuclear envelope, promoting the re-organization of PI(4,5)P2 and the coupling of the plasma membrane Ca^2+^ channel Orai1 with the ER Ca^2+^ sensor STIM1 ^63^. Recent work shows that septins corral PI(4,5)P2 in membranes ^64^ and synergize with Cdc42, WASP/WAVE and Arp2/3 in remodeling actin into a ring-like structure at Ora1-STIM1 contact sites ^65^. Interestingly, ER-plasma membrane contact sites were recently found to regulate focal adhesion turnover through lipid exchange in response to Ca^+2^ influx ^66^. Therefore, SEPT9_i1 may promote the nucleation of invadopodia at contact sites of the nuclear envelope with the plasma membrane. In this scenario, nuclear envelope folds and indentations would diminish contacts with the plasma membrane. Furthermore, nuclear deformities may hinder actin polymerization at the interface of the nucleus with the plasma membrane. As polymerization of invadopodial actin exerts forces against the plasma and nuclear membranes, an unstable nuclear envelope may not be able to support the elongation and protrusion of invadopodia into the ECM ^23, 67^.

In endothelial cells, septins are posited to provide a diffusion barrier downstream for the maturation of podosomes by forming rings on phosphatidylinositol-rich domains that corral β-integrin ^40^. Septin ring assembly is temporally downstream of Tks5 clustering and actin nucleation ^40^. In breast cancer cells, however, septins appear to function upstream of invadopodia precursor clustering, because the latter is diminished by septin depletion. In the early stages of invadopodia formation, we envisage that septins act at the interface of the nuclear envelope and plasma membrane, interacting with the machinery of actin nucleation. Lack of an invadopodia phenotype in cells depleted of SEPT9_i2, an isoform of diminished association with actin ^33^, suggests that SEPT9_i1 functions through actin. If SEPT9_i1 is a component of the LINC complex, it may scaffold the nuclear bound FHOD formins and/or crosslink perinuclear actin with cell-ECM adhesions. Interestingly, septins regulate FHOD1 phosphorylation at sites of bacterial entry ^68^ and SEPT12, a testis-specific septin of the SEPT3 group of septins which includes SEPT9, interacts with a SUN domain protein (sperm-associate antigen 4) and lamin B1, influencing the localization of these proteins ^69, 70^. Thus, septins may enable a crosstalk between the actin assembly machinery of the nuclear and plasma membranes, favoring the formation of invadopodia at juxtanuclear sites of the plasma membrane. This raises the possibility that juxtanuclear invadopodia are mechanistically and qualitatively distinct from invadopodia in other subcellular regions, differing in stability, protrusive and/or proteolytic activity.

Septins filaments localize preferentially on juxtanuclear regions of the cytoplasm in various cell types, but it has been unknown whether this is due to interactions with the nuclear envelope. Here, we have fortuitously discovered that SEPT9_i1 influences nuclear shape and integrity. SEPT9_i1 depletion caused nuclear envelope deformities such as nuclear grooves and indentations, and altered nuclear shape. Rescue of nuclear shapes by lamin A indicates that SEPT9_i1 contributes to the mechanical rigidity of the nucleus, which in turn might be necessary for the mechanical support and outgrowth of invadopodia. Septins are known enhancers of cortical membrane stiffness and can function similarly on the nuclear envelope ^71, 72^. We hypothesize that SEPT9_i1 promotes nuclear rigidity from the cytoplasmic side of the nuclear envelope, where it could act synergistically with actomyosin in enhancing or maintaining membrane tension. Loss of the latter would cause nuclear envelope wrinkles and invaginations, consistent with the observed phenotypes ^73, 74^. We cannot exclude, however, the possibility of SEPT9_i1 functioning in the nuclear lamina. The N-terminus of SEPT9_i1 contains a unique amino sequence with a bipartite nuclear localization signal, which interacts with importin α ^62^, and SEPT9 has been previously observed in the nucleolus ^27^.

Throughout our studies, knock-down of SEPT9_i2 had no to little effect on invadopodia and nuclear shape, underscoring the isoform-specific functions of the oncogenic SEPT9_i1. This correlated with a scarcity of perinuclear SEPT9_i2 filaments, as also observed in previous work which proposed that SEPT9_i2 has tumor suppressing properties inhibiting cell migration ^33^. The SEPT9_i1 isoform, therefore, may interact with the nuclear envelope through its unique N-terminal sequence that binds importin α ^62^. The N-terminus of SEPT9_i1 also interacts with Ruk/CIN85 (regulator of ubiquitous kinase/Cbl-interacting protein of 85 kDa), an EGF receptor (EGFR) adaptor protein, which regulates the degradation and trafficking of EGFR ^75, 76^. Because CIN85 localizes to EGFR-carrying endosomes ^77^, perinuclear SEPT9_i1 levels may increase in response to EGF due to association with perinuclear endosomes. Recent studies have shown that endosomes are captured by the nuclear envelope in response to receptor tyrosine kinase activation ^78^. Nuclear envelope-anchoring of MMP-containing endosomes is critical for degrading ECM at perinuclear regions ^21^.

We posit that SEPT9_i1 functions as a downstream effector of EGFR signaling, which biases the formation of invadopodia at juxtanuclear sites of the plasma membrane. Strategically, this would facilitate nuclear threading through ECMs of low penetrability by focusing ECM degradation around the nucleus, which is the largest physical impediment in cell translocation. It is plausible that in addition to EGF, mechanotransducing signals from highly dense ECMs upregulate septin localization at the nuclear membrane, which in turn enhances nuclear envelope rigidity and the formation of juxtanuclear invadopodia. In support of this possibility, ECMs of higher stiffness upregulate the expression of Cdc42EP3 which promotes septin filament formation and association with actin in cancer-associated fibroblasts ^79–81^. Stiff ECM environments are also linked to septin upregulation in the context of epithelial-to-mesenchymal transition ^82^, and SEPT9_i1 promotes the invasive strategies of melanoma cells through collagen-rich ECMs ^83^. Thus, SEPT9_i1 might be part of an adaptive response in cancer cells that cannot translocate through dense ECMs in spite of having deformable nuclei.

Our findings have important implications for the function of SEPT9 isoforms in the pathogenesis of breast cancer metastasis. First, the impact of SEPT9 on nuclear morphology and properties raises the possibility of genomic and/or epigenomic alterations owing to the association of chromatin with the nuclear envelope. Indeed, over-expression of SEPT9 isoforms 1, 2 and 3 have differential effects on the transcriptome of breast cancer MCF-7 cells ^36^, and isoform 4 reduces the size of the nucleus ^84^. Alternatively, changes in nuclear morphology may impact the nucleocytoplasmic shuttling of transcriptional regulators such as YAP. Of note, SEPT2 depletion reduces the nuclear translocation of YAP in response to ECM of higher stiffness ^79^. Second, variations in SEPT9 isoform expression are likely to influence the invasive strategies of cancer cells, which in the long-term can be of both diagnostic and therapeutic value. In light of recent evidence showing that lamin A/C is downregulated in breast cancer cells ^85^, amplification of SEPT9 could shift some breast cancers to an invadopodia-based invasion. Third, SEPT9 isoforms may impact the genomic instability of breast cancers during metastatic migration. By reducing nuclear deformability, SEPT9_i1 may prevent further DNA damage and genomic alterations that could alter the highly invasive phenotype of metastatic cancer cells. Lastly, our results indicate that SEPT9_i1 localization and function in the formation of juxtanuclear invadopodia is upregulated by EGF signaling, and thus SEPT9 might be part of the malignant programming of EGFR-positive breast cancers. Collectively, our findings open new avenues of research into the function of septins in nuclear mechanobiology and cancer cell invasion, which in the long-term could benefit treatments of metastatic cancer.

### Limitations of the study

Our studies were technically limited to use of tissue culture cell lines and in vitro assays of ECM degradation and invasion, lacking evidence from an in vivo environment. Intravital imaging of metastatic cancer cells and/or ex vivo 3D organoid models are necessary for probing septin functions in invadopodia formation and the mechanical properties of the nucleus. Investigation of the latter can also benefit from approaches that utilize microfabricated microchannels, 3D collagen gels, atomic force microscopy and biosensors of mechanical forces. Conceptually, our study is limited by lack of understanding of how septins are precisely linked to the nuclear envelope and the nuclear-associated mechanisms of actin nucleation and elongation, which may feedback to invadopodia formation at the interface of the nuclear and plasma membranes. This work did not address septin roles in invadopodia elongation and MMP secretion, which require tools that decouple temporally and functionally septins from upstream mechanisms of invadopodia formation.

## STAR Methods

### Key resources table

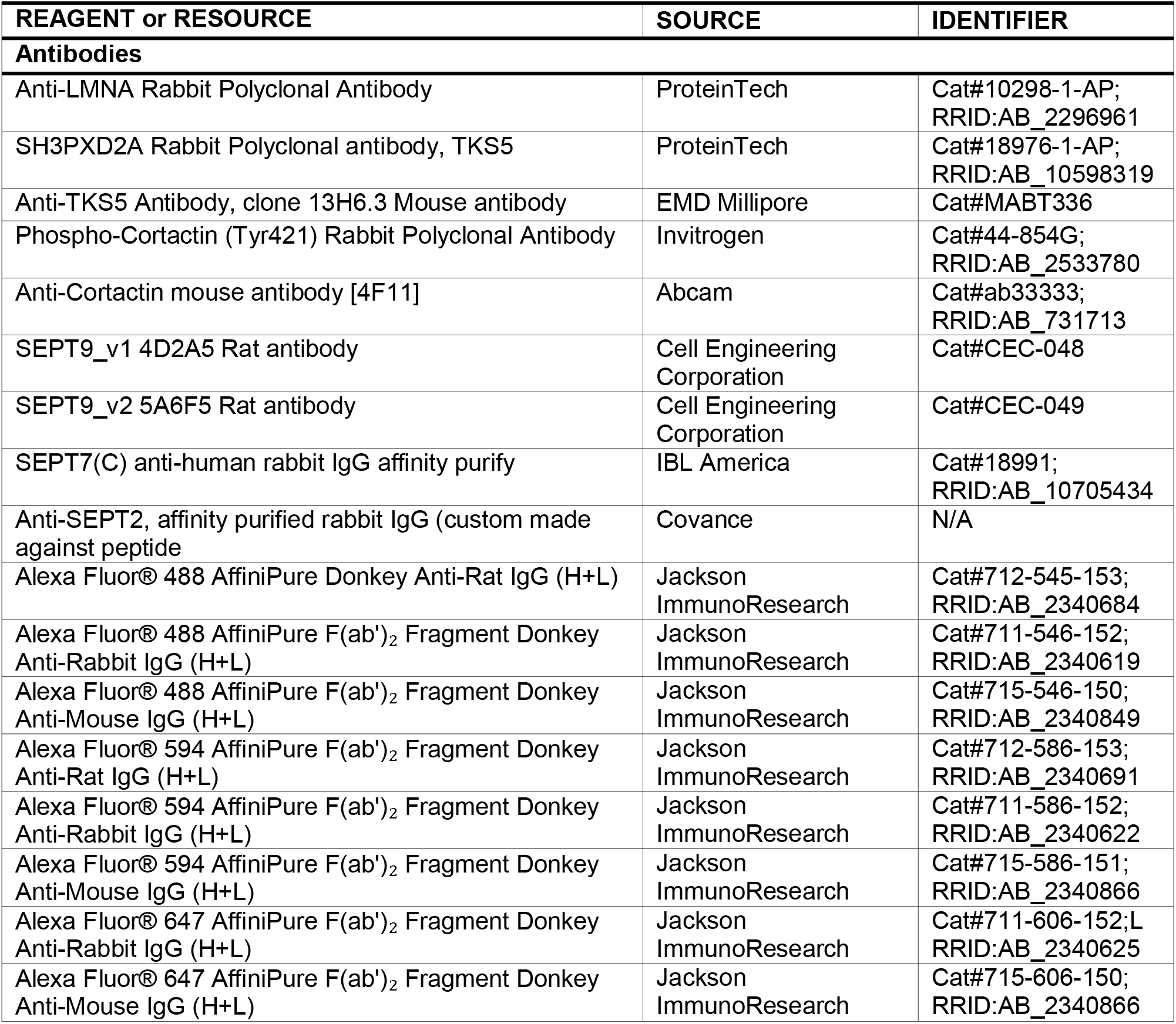
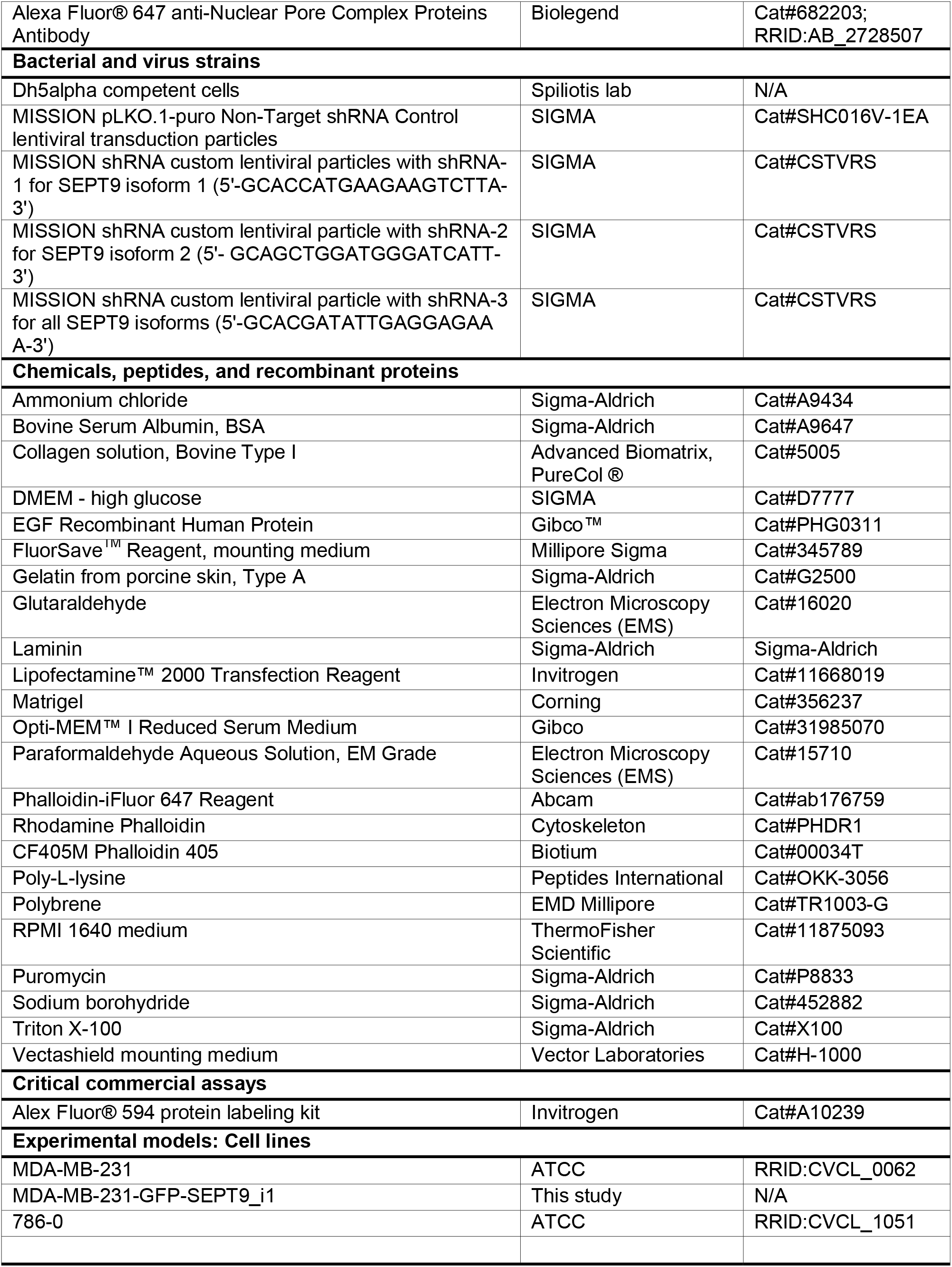
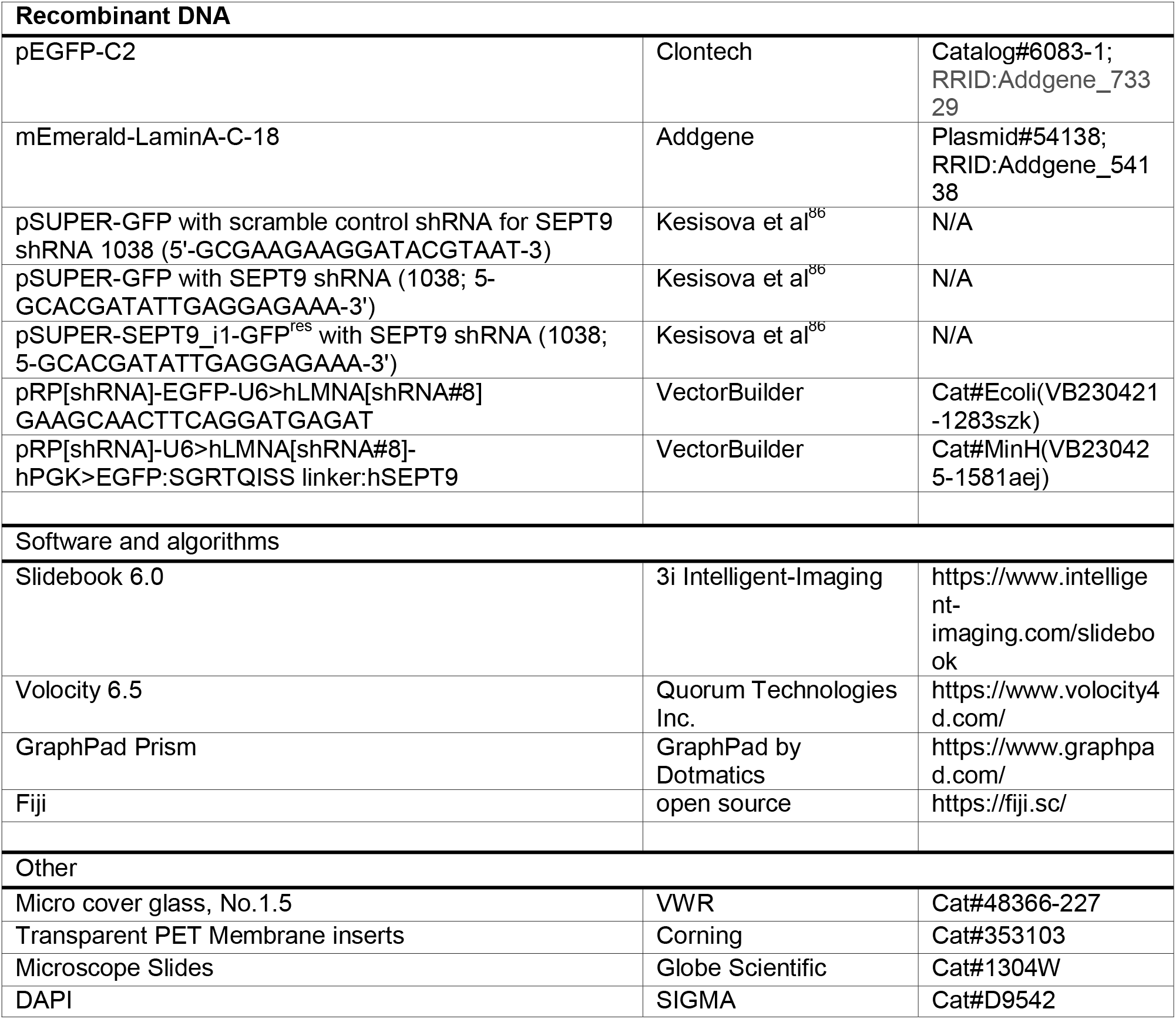

### Resource Availability

#### Lead Contact

Requests for further information, resources and/or reagents should be made to the lead contact: Dr. Elias Spiliotis

#### Materials availability

Reagents generated in this study and are not available at Addgene can be made available upon request, but may require payment and/or a material transfer agreement.

### Experimental model and subject details

#### Cell lines

MDA-MB-231 (ATCC HTB-26) and MDA-MB-231 cells that stably express GFP-SEPT9_i1 were maintained in DMEM High Glucose medium containing 1g/L sodium bicarbonate and supplemented with 10% heat-inactivated FBS, and 1% PSK (penicillin, streptomycin, kanamycin). 786-0 (ATCC CRL1932) cells were maintained in RPMI 1640 Medium supplemented with 10% FBS and 1% PSK. Cells were maintained at 37°C in 5% CO_2_. MDA-MB-231 cells that stably expressed GFP-SEPT9_i1 were generated after transfection and selection with G418.

### Method Details

#### Cell transductions by lentiviral shRNAs and transfections with plasmids

Using the SIGMA Custom Mission shRNA Lentiviral service, lentiviruses were made to order (SIGMA # CSTVRS) with the following shRNA sequences inserted into the pLKO.1puro vector: GCACCATGAAGAAGTCTTA (shRNA1), GCAGCTGGATGGGATCATT (shRNA2) and GCACGATATTGAGGAGAAA (shRNA3). ShRNAs 1 and 2 targeted unique sequences in human SEPT9 isoforms 1 and 2, respectively, and shRNA 3 targeted a sequence shared by all SEPT9 isoforms. MISSION pLKO.1-puro non-target shRNA control transduction particles (SIGMA, #SHC016V) were used as controls. MDA-MB-231 cells (5×10^5^) were seeded one day prior to transduction with a viral mixture - lentiviruses with shRNA sequences at a 2:1 ratio of viral particle to cell, and 8 µg/mL Polybrene (EMD Millipore, Cat #TR1003-G). Cells were incubated with viral mixture for 24 h after which the growth medium was replaced. After another 24 h incubation, cells were cultured with media containing puromycin (2 µg/ml) for 48-72 h before trypsinization and plating for use in experimental assays. For 2D gelatin degradation assays (dark or fluorescently-labelled gelatin), cells were seeded directly onto gelatin-coated coverslips, allowed to adhere for 1 h and then treated with 10 nM EGF for 4h, fixed and immunostained. For 3D chemoinvasion assays, transduced cells were seeded on Matrigel- and laminin coated Transwells, and incubated for 12-24 h. For rescue experiments with mEmerald-lamin A, cells were transfected with plasmid DNA after 48 h of transduction with shRNA-containing lentivirus, and seeded on gelatin 24 h following transfection of plasmid DNA. All seeded cells were incubated to adhere for 1 h and then treated with EGF for 4 h.

MDA-MB-231 and 786-0 cells were transfected with pSUPER plasmids encoding for GFP or shRNA-resistant SEPT9_i1^res^-GFP and human SEPT9 shRNA (5’-GCACGATATTGAGGAGAAA-3’) or scramble control shRNA (5’-GCGAAGAAGGATACGTAAT-3’). These plasmids were constructed and described previously ^86^. The plasmid pRP[shRNA]-EGFP-U6>hLMNA[shRNA#8] encoding for GFP and a human lamin A shRNA targeting the sequence 5’-GAAGCAACTTCAGGATGAGAT-3 was purchased from VectorBuilder (catalog# Ecoli(VB230421-1283szk)). For rescue of lamin A depletion with SEPT9_i1, the same plasmid was modified to add the sequence of SEPT9 to the C-terminus of GFP using the linker SGRTQISS. The new plasmid pRP[shRNA]-U6>hLMNA[shRNA#8]-hPGK>EGFP:SGRTQISS linker:hSEPT9[NM_001113491.2] was designed and purchased from VectroBuilder (catalog# MinH (VB230425-1581aej).

#### Cell transfections with plasmid DNA

Following a 48 h transduction with lentiviruses as described above, cells were transfected with mEmerald-Lamin A-C-18 (addgene plasmid # 54138) and pEGFP-C2 (Clontech) by initially combining plasmid DNA with the Lipofectamine™ 2000 Transfection Reagent (Invitrogen, Cat#11668019) in Opti-MEM™ I Reduced Serum Medium (Gibco, Cat#31985070) for 20 minutes, and diluting the mix in cell media. The culture media of transduced cells were replaced with the plasmid DNA-containing media for 6 h, after which there was a change and overnight incubation with fresh media. Subsequently, cells were dissociated and plated on gelatin-coated coverslips as described below. All other transient transfections with shRNA-encoding plasmids were performed by combining plasmid DNA with the Lipofectamine™ 2000 Transfection Reagent (Invitrogen, Cat# 11668019) in Opti-MEM™ I Reduced Serum Medium (Gibco, Cat# 31985070) for 20 minutes, and adding the mix dropwise into cell culture media. Following 4-6 h incubation, transfected cell cultures were replaced with fresh complete growth medium and allowed to incubate for up to 72hrs before seeding on matrix-coated coverslips for immunostaining.

#### Gelatin degradation assays

Gelatin was labeled using Alex Fluor® 594 protein labeling kit (Invitrogen, Cat#A10239) according to the manufacturer’s instructions. For fluorescently-labeled gelatin coating, sterilized no. 1.5 glass coverslips (VWR, Cat # 48366-227) were incubated in 2 ml of ice-cold 50 µg/mL poly-l-lysine (Peptides International, Cat # OKK-3056) for 20 minutes. Following incubation, coverslips were rinsed three times in PBS (filter sterilized) and then incubated in 2 ml of 0.5% Glutaraldehyde (Electron Microscopy Sciences, Cat # 16020) for 15 minutes at room temperature. Coverslips were then rinsed and left to incubate in PBS as gelatin deposits were prepared. Unlabeled 2% gelatin (Sigma, Cat# G2500) was mixed with Alexa Fluor 594-labeled gelatin at 8:1 ratio after allowing gelatin to fully dissolve in a 37°C water bath. A drop (100 µl) of fluorescent gelatin mix was deposited on a flat parafilm, and coverslip was inverted onto the drop with its poly-l-lysine- and glutaraldehyde-coated surface facing down, and incubated for 10 minutes in the dark. For dark coverslip coating with dark non-fluorescent gelatin, 2% unlabeled gelatin was deposited on a parafilm. Following incubation, gelatin-coated coverslips were rinsed in PBS and quenched with 5 mg/ml sodium borohydride (Sigma, Cat # 452882) for 15 minutes at room temperature. Gelatin-coated coverslips were then rinsed in PBS extensively to remove bubbles and rinsed twice with cell culture media before cell plating.

MDA-MB-231 cells (3 x 10^4^) and 786-0 cells (3 x 10^4^) were diluted in 2 ml of cell culture media and seeded onto a gelatin-coated coverslip. For EGF treatments, 10 nM of EGF (Gibco™ Cat#PHG0311) was added to the growth medium after 1 h of plating cells, and then incubated for 4 h – total incubation time of 5 h. Following incubation, cells were fixed for 15 minutes in 4% PFA (Electron Microscopy Sciences, Cat # 15710) and 4% sucrose (Sigma, Cat # S9378) in PBS and immunostained.

#### Transwell chemoinvasion assays

Transparent polyester (PET) membrane Transwell inserts with 1.0 µm pores (Corning, Cat # 353103) were coated with 100µL of a matrix cocktail – 10 mg/mL Matrigel (Corning, Cat # 356237) and 0.5 mg/mL laminin (Sigma Cat # L2020) – and incubated at 4°C with gently rocking for 1 h. Matrix mix (90µL) was removed and inserts were incubated at 37°C for 30 minutes. Subsequently, inserts with polymerized matrix were hydrated on both sides with DMEM High Glucose medium supplemented with 10% FBS and 1% PSK (Penicillin, Streptomycin and Kanamycin). EGF (10 nM; Gibco Cat # PHG0311) was added to the media of the bottom chamber of the Transwell insert in order to create a chemo-attractive gradient. Cells (3 x 10^4^) were resuspended in 1 ml of complete growth medium and added to the top side of the Transwell insert and incubated for 12 or 24 h. Transwells with cells and EGF chemoattractant were incubated for 12hr and 24hr. This protocol was adapted from previous work by the Vignjevic group ^87^.

#### Immunofluorescence

On gelatin-coated glass coverslips, cells were fixed for 15 minutes with 4% paraformaldehyde (Electron Microscopy Sciences, Cat # 15710) in PBS or 4% PFA (Electron Microscopy Sciences, Cat#15710) and 4% sucrose (Sigma, Cat # S9378) in PBS if the coverslips were coated with fluorescent gelatin. Following fixation, coverslips were rinsed with PBS and quenched with 0.4% ammonium chloride (Sigma, Cat # A9434) in PBS. Cells were permeabilized with 0.1% Triton X-100 (Sigma Cat # X100) for 15 minutes and blocked with 2% Bovine Serum Albumin (BSA; Sigma, Cat # A9647) for 1 at room temperature. Coverslips were washed twice in PBS following blocking step, and incubated with primary antibody overnight at 4°C, and then incubated with secondary antibody for 1 h at room temperature. Immunostained coverslips were mounted with FluorSave™ Reagent (Millipore, Cat # 345789) onto frosted microscope slides (Globe Scientific, Cat # 1304W).

Cells on Transwell membrane inserts were rinsed in warm PBS and fixed for 15 minutes with 4% PFA (Electron Microscopy Sciences, Cat # 15710) in PHEM buffer (60 mM PIPES, 25 mM HEPES, 10 mM EGTA, 4Mm MgSO_4_, pH 6.9). Fixed inserts were rinsed three times in PBS, quenched with 0.5% NH_4_Cl (Sigma Cat # A9434) and again rinsed three times in PBS. For immunostaining, inserts were incubated with 0.1% Triton X-100 (Sigma Cat # X100) for 15 minutes and blocked with 2% Bovine Serum Albumin (BSA; Sigma Cat # A9647) for 1 h at room temperature. Both sides of insert were incubated with primary antibodies overnight and secondary antibodies for 1 h with PBS rinses in between each step. Subsequently, Transwell membranes were gently cut out of inserts with a scalpel razor blade and placed on microscope slides (Globe Scientific, Cat # 1304W) with their apical side facing up. Upon adding a drop of Vectashield mounting medium (Vector Laboratories, Cat # H-1000), a glass coverslip (VWR, Cat # 48366-227) was placed on the membrane and the edges were sealed with clear nail polish after dabbing excess of mounting media from the edges of the coverslip. Mounted membranes were left to dry in the dark for more than 1 h at room temperature. Slides were imaged once nail polish was completely dry.

Cells that were plated on collagen were stained as described above for cells plated on gelatin. Prior to plating, 22mm x 22mm no. 1.5 glass coverslips (VWR, Cat#48366-227) were coated with PureCol Type I Bovine collagen (Advanced Biomatrix, Cat # 5005) at 10 µg/cm^2^ for 10 minutes, then allowed to air-dry for 30 minutes before placing them under UV for another 30 minutes.

The SEPT9 isoforms SEPT9_i1 and SEPT9_i2 were stained with isoform-specific rat monoclonal antibodies 4D2A5 (Cell Engineering Corporation; Cat # CEC-048) and 5A6F5 (Cell Engineering Corporation; Cat # CEC-049) ^33^, respectively, which were generously gifted by Dr. Taro Tachibana (Cell Engineering Corporation, Japan). Rabbit polyclonal antibodies were used for staining lamin A/C (Proteintech, Cat # 10298-1-AP), TKS5/SH3PXD2A (Proteintech, Cat # 10298-1-AP), phospho-cortactin (Tyr2421; Invitrogen, Cat#44-854G), SEPT7 (IBL America, Cat # 18991) and a custom-made affinity purified antibody against SEPT2, which was generated by Covance using the antigenic peptide MSKQQPTQFINPETPGYVC. Mouse monoclonal antibodies were used for staining TKS5 (clone 13H6.3; EMD Millipore, Cat # MABT336-25UG) and cortactin (clone 4F11; Abcam Cat # ab33333). Nucleoporins were labeled with Alexa Fluor® 647 anti-Nuclear Pore Complex Proteins Antibody (mAb 414; Biolegend Cat # 682203). Actin was labeled with CF405M phalloidin 405 (Biotium Cat # 00034T), rhodamine phalloidin (Cytoskeleton Cat # PHDR1) or phalloidin-iFluor 647 reagent (Abcam ab176759). Nuclei were labeled with DAPI (Sigma Cat # D9542).

The following secondary antibodies were used: Alexa Fluor® 488 AffiniPure Donkey Anti-Rat IgG (H+L) (Jackson ImmunoResearch, Cat # 712-545-153), Alexa Fluor® 488 AffiniPure F(ab’)₂ Fragment Donkey Anti-Rabbit IgG (H+L) (Jackson ImmunoResearch, Cat # 711-546-152), Alexa Fluor® 488 AffiniPure F(ab’)₂ Fragment Donkey Anti-Mouse IgG (H+L) (Jackson ImmunoResearch, Cat # 715-546-150), Alexa Fluor® 594 AffiniPure F(ab’)₂ Fragment Donkey Anti-Rat IgG (H+L) (Jackson ImmunoResearch, Cat # 712-586-153), Alexa Fluor® 594 AffiniPure F(ab’)₂ Fragment Donkey Anti-Rabbit IgG (H+L) (Jackson ImmunoResearch, Cat # 711-586-152), Alexa Fluor® 594 AffiniPure F(ab’)₂ Fragment Donkey Anti-Mouse IgG (H+L) (Jackson ImmunoResearch, Cat # 715-586-151), Alexa Fluor® 647 AffiniPure F(ab’)₂ Fragment Donkey Anti-Rabbit IgG (H+L) (Jackson ImmunoResearch, Cat # 711-606-152), and Alexa Fluor® 647 AffiniPure F(ab’)₂ Fragment Donkey Anti-Mouse IgG (H+L) (Jackson ImmunoResearch, Cat # 715-606-150).

#### Microscopy

Wide-field imaging was performed with the Zeiss AxioObserver Z1 inverted microscope equipped with a Zeiss 63x/1.4 NA oil objective and a Hamamatsu Orca-R2 CCD camera (Figures 1A-B, 2A-B, 3A, 6A, S1). Images acquired using Slidebook 6.0 acquisition software. Z-stacks were acquired with a 0.27 µm step-size in the z-direction. Images were processed after acquisition, where indicated, with Slidebook 6.0 constrained iterative or nearest neighbor deconvolution module.

Laser scanning confocal microscopy was performed with a Leica STELLARIS microscope using a 63X/1.4 NA oil objective, Power HyD detectors and a zoom in of 1.28 F (Figures 4A-B, 4D-G, 5C-F, 6D). The LIGHTNING deconvolution module was used to generate super-resolution images from 3D stacks. Alternatively, an Olympus FV1000 microscope was used with a 60X/1.42 NA oil objective (Figures 1D-G, 5A-B and S3A-D).

Spinning disk confocal microscopy (Figure 4C) was performed on an inverted Olympus IX83 microscope equipped with a Yokogawa SCU-10 spinning disk scanner, a 60X/1.49 NA silicone oil objective, a Hamamatsu digital CMOS camera Orca Flash 4.0 V2 and 405/488/561/635 nm excitation lasers. All z-stacks were taken at ∼0.2 µm step intervals in the z-direction.

#### Image quantification, processing and analysis

In Figure 1, analysis of SEPT9_i1 localization at areas of gelatin degradation was done manually in Fiji. Areas of gelatin degradation were masked using manual thresholding of inverted gelatin channel. Particle analysis was performed on the gelatin masks in order to acquire ROIs with precise areas of degradation. In these ROIs, fluorescence intensity of SEPT9_i1 was measured for each cell. Recorded values were categorized into ROIs with surface area of < 4 µm^2^, 4-8 µm^2^ and > 8 µm^2^, and then imported into GraphPad Prism for graphing and statistical analysis. SEPT9_i1 localization in 3D invadopodia was also quantified in Fiji by using the rectangle tool to mark the full length of invadopodia in the z-axis, and divide it into three segments of equal length, which were proximal, medial and distal with respect to the cell body. Presence or absence of SEPT9_i1 was scored for each segment of the invadopodium. The number of invadopodia that contained SEPT9_i1 in each of these segments was quantified as percentage of total invadopodia. Data were imported into GraphPad Prism for statistical analysis and graphing.

In Figure 2, depletion of SEPT9 in cells was quantified in Fiji by measuring the average intensity values of SEPT9 per cell surface area. Intensity measurements for each image was normalized to background of the corresponding image, by manual background subtraction – intensity of background area is subtracted from intensity of cell area. Cell surface area were masked by boosting the fluorescence signal of SEPT9 to levels that cell edges were visible, and cell contours were drawn. Quantification of the areas of gelatin degradation was performed with the Slidebook 6.0 software using the “segment mask” tool to generate masks of the total cell area (Mask Cell). Using the cellular mask outline, a mask of the same area was made in the fluorescence channel of gelatin (Mask Gelatin). Using the “mask operations” tool, the Mask Gelatin was subtracted from the Mask Cell to generate a mask (Mask Degradation) with the degraded gelatin area per cell surface. In Figure S6C, masks of nuclear and gelatin degradation areas were created as above, and the total surface area of degraded gelatin that overlapped with the nuclear region was divided by the surface area of the nucleus. The ratios were multiplied by 100 to derive the percentage of gelatin degraded underneath the nucleus. Values were exported into Excel prior to entry into GraphPad Prism for statistical analysis and graphing.

In Figures 2G-H and S2F, quantification of invadopodia number and length in 3D chemoinvasion assays was done in Fiji. Invadopodia were identified as phalloidin-stained structures that protruded into basal side of the porous Transwell membrane, and counted within the surface area of their respective cell bodies, which were outlined based on the phalloidin stain or GFP fill on the apical side of the Transwell membrane. Invadopodia lengths were measured in 3D cross-section images using the line tool.

In Figures 3, 5J and S6E, quantification of the number and size of ventral clusters of TKS5, cortactin and phosphor-Cortactin (pY421) was performed in Fiji. Maximal intensity projections were generated from ventral optical sections that span a z-axis distance of ∼1 µm from the basal cell surface. Using the Particle Analysis, puncta were segmented as ROIs using ranges for fluorescence intensity, pixel size area (>0.22 μm^2^) and roundness values (0.0 to 1.0), which most accurately masked, identified and distinguished clusters from background staining. Segmentation thresholds for a protein of interest were set in control cells and subsequently applied uniformly to all other conditions. Number of puncta per cell and average size of puncta per cell were quantified in Fiji, and data were statistically analyzed and plotted in Prism GraphPad.

In Figure 4C, the fluorescence intensity plot was made using the straight line tool in Fiji. A 2 µm-thick line was drawn across a section of the nuclear envelope (Figure 4B, inset). Grey value data for the fluorescence intensities of SEPT9 (green) and nucleoporins (magenta) were exported in excel and normalized to the largest intensity value of each fluorescence channel. Plot shows normalized intensity values.

In Figures 5 and S4F, morphometric quantifications of the nucleus were performed in Fiji. Perimeter lengths and surface areas were automatically derived from nuclei, which were masked as ROIs using the free hand tool to trace the edges of lamin A/C and nucleoporin fluorescence in images of maximum intensity projection. The complexity index was calculated as the ratio of perimeter length to surface area. For volume analysis, nuclei in each optical section of a z-stack were outlined and marked as individual ROIs. The surface of the ROI of each optical slice was measured and multiplied by the step size to derived the volume in between successive optical sections. The net volume of each nucleus was derived by adding all the volumes of all optical section across the z-axis of the nucleus. In Figure 6H, nuclear grooves were quantified in 3D images of nucleoporin-stained nuclei. Using the nucleoporin fluorescence channel, lines of nucleoporin that extended inward from the nuclear rim were counted in all optical slices across the z-axis of a nucleus and summed to derive the total number of nuclear grooves per cell.

In Figure S3E, lamin A/C quantifications were performed from 3D image stacks of confocal microscopy. For each nucleus, every single optical slice was segmented along the entire z-axis of the nucleus to generate a region of interest (ROI) in Fiji. The mean fluorescence intensity was derived for each optical slice of the nucleus and values were averaged to obtain the mean fluorescence intensity per nucleus.

In Figures 6B, S5F and S6F, quantifications of percentage of juxtanuclear puncta with TKS5, actin and phospho-cortactin (pY421) were performed by generating maximal intensity projections of the optical section images of the ventralmost ∼1 µm of the cell height. In these maximal projections, the DAPI channel was used to generate a mask that marks the footprint of the nucleus. Puncta with TKS5, actin and cortactin-pY421, which overlapped with the nuclear footprint and a 3 μm-wide zone around the nuclear footprint edge were marked manually, and quantified as percentage of total puncta per cell (percentage of juxtanuclear clusters). As above, the Particle Analysis tool was used. Puncta were segmented as ROIs using ranges for fluorescence intensity, pixel size area (>0.22 μm^2^) and roundness values (0.0 to 1.0), which most accurately masked, identified and distinguished clusters from background staining. Segmentation thresholds for a protein of interest were set in control cells and subsequently applied uniformly to all other conditions. Percentage of juxtanuclear puncta was recorded per each cell and plotted in Prism GraphPad.

In Figure 6C, quantification of the juxtanuclear levels of SEPT9_i1 was performed in ventralmost optical sections, which contained a mask of the nuclear footprint area from maximum intensity projections of the nucleoporin signal. Using Fiji, fluorescence segmentation and mask subtractions, we generated masks of the SEPT9_i1 fluorescence signal that overlapped with the nuclear footprint. The surface area of the latter mask was divided by the surface area of the nucleus to derive the fraction of the nuclear area occupied by SEPT9_i1. The fraction number was multiplied by 100 to derive the percentage of the nuclear area overlapping with SEPT9_i1.

In Figure 6E, the total number of puncta containing actin and cortactin was quantified in maximum projections of sections which were acquired from the ventralmost 1 μm of each cell using fluorescence thresholding and a 0.22 μm^2^ size cutoff. The number of juxtanuclear puncta was quantified as percentage of total puncta as performed in the quantifications of Figure 6B.

Three-dimensional rendering of confocal microscopy images was performed in the Volocity 6.5 software (Figures 1D-G, 4F-G, 5B) using the “fine filter” method for all channels or in the 3D viewer of the Leica STELLARIS software. Three-dimensional stacks of wide-field images were deconvolved with constrained iterative (Figure 3) or nearest neighbor deconvolution (Figure 6) in the Slidebook 6.0 software.

### Statistical Analyses

Statistical analyses were performed with the Prism GraphPad software. Depending on their n values, all data sets were initially subjected to normality tests (Kolmogorov-Smirnov, Shapiro-Wilk, D’Agostino & Pearson) to assess whether they have a normal Gaussian or non-normal distribution. Descriptive statistics were also used to determine whether data sets have equal or unequal standard deviations (SDs). Pair-wise comparisons of data with Gaussian distributions were analyzed with a parametric unpaired t-test. A Welch’s correction was applied if SDs were unequal. Pair-wise comparisons of data with non-normal distributions were analyzed with the non-parametric Mann-Whitney test. More than two groups of data were analyzed with one-way ANOVA. If all data sets were normally distributed with equal SDs, an ordinary ANOVA was used with a post-hoc Dunnett or Bonferroni test for multiple pair-wise comparisons. For normally distributed data with unequal SDs, a Welch ANOVA test was performed with a post-hoc Dunnett T3 test for multiple comparisons. For data that were not normally distributed, a non-parametric Kruskal-Wallis ANOVA test was performed with a post-hoc Dunn’s test for pair-wise comparisons. A chi-squared test was applied for data that were binned into categorical groups. Results with p values of < 0.05 were considered to be statistically significant.

## Data and code availability

- All data reported in this paper will be shared by the lead contact upon request
- This paper does not report original code
- Any additional information required to reanalyze the data reported in this paper is available from the lead contact

## Acknowledgments

We thank Dr. Ryan Petrie (Drexel University) and his laboratory for generous help with reagents and technical advice, and Dr. Taro Tachibana (Cell Engineering Corporation, Osaka, Japan) for his generous gift of SEPT9 isoform 1 and 2 antibodies. We are also grateful to Dr. Harini Sreenivasappa (Drexel University) for help with microscopy. All microscopy was performed at Drexel University’s Cell Imaging Center. The work was supported by National Institutes of Health grant 5R35 GM136337-03 (ETS), a Commonwealth Universal Research Enhancement Formula (CURE) grant SAP 4100079710 from the Pennsylvania Department of Health (ETS), and Department of Defense Breakthrough Award W81XWH-19-1-0104 (CM).

## Author Contributions

J.O. designed and conducted experiments, analyzed data and wrote sections of the manuscript. D.A. developed assays, performed experiments and analyzed data. T.M.J. conducted experiments and analyzed data. C.M. developed cell lines and contributed to the writing of the manuscript. E.T.S. procured funding, directed the design and progress of all experiments and analyses, acquired data, assembled figures and wrote the manuscript.

## Inclusion and Diversity

We worked to ensure diversity in experimental samples through the selection of the cell lines. One or more authors of this paper identifies as an underrepresented ethnic minority in their field of research or within their geographical location. One or more authors of this study identifies as a gender minority in their field of research. While citing references scientific relevant for this work, we also actively worked to ensure gender balance in our reference list.

