## Supplemental Figures for "An oncogenic isoform of septin 9 promotes the formation of juxtanuclear invadopodia by reducing nuclear deformability"

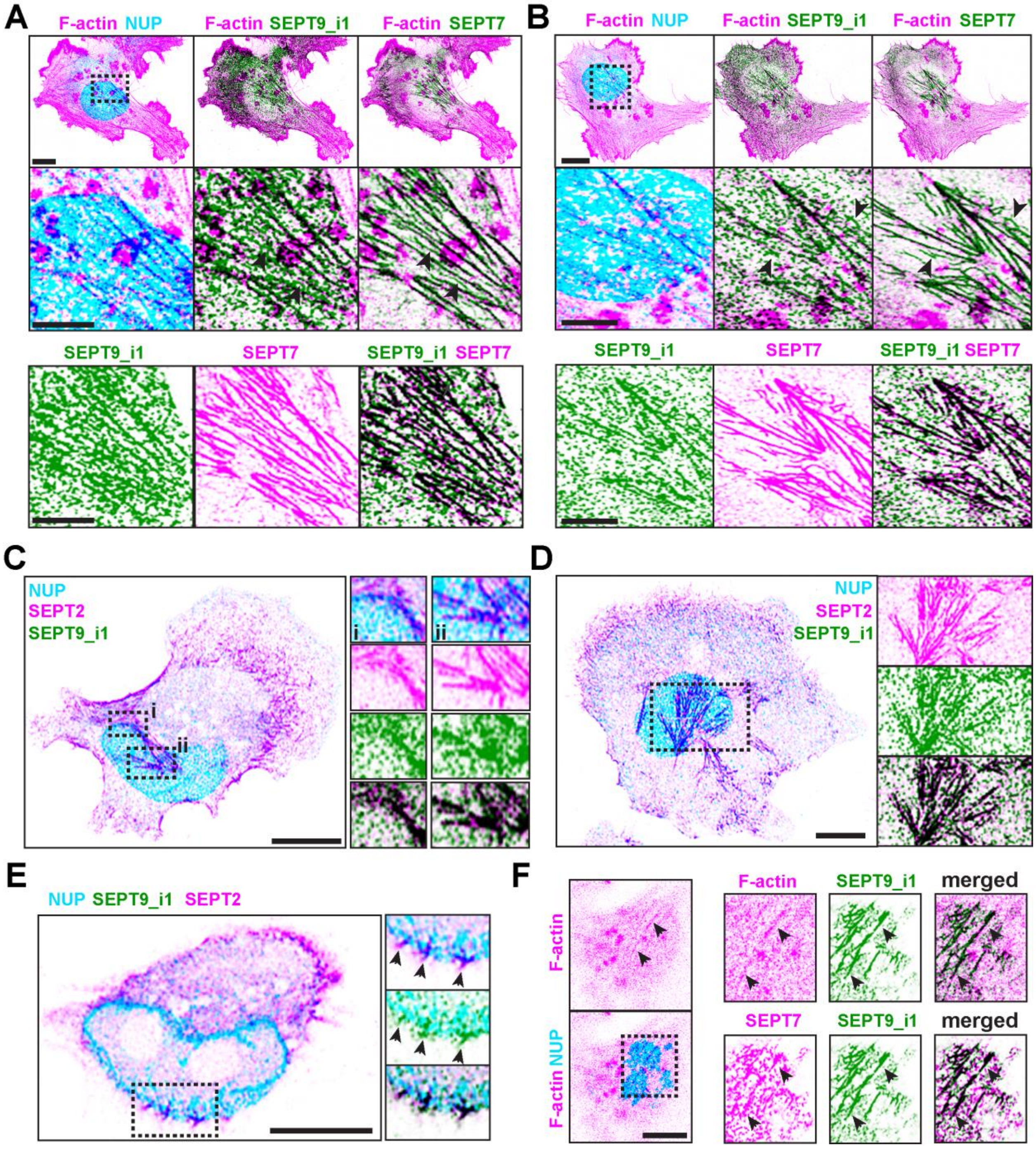


**Figure S1. SEPT9_i1 colocalizes with SEPT7 and SEPT2, related to Figure 1.**

(A-B) Super-resolution confocal microscopy images show ventral optical sections from MDA-MB-231 cells, which were plated on gelatin (A) or collagen (B) for 5 h in the presence of EGF prior to staining with phalloidin (F-actin) and antibodies against SEPT7, SEPT9_i1 and nucleoporins (NUP). Septin and actin localizations are shown in higher magnification for the outlined region (dashed rectangle) in the middle and bottom panels. Black arrowheads point to septin filaments that do not colocalize with actin filaments. Overlaying regions between inverted magenta and green colors are shown in black. Scale bars, 10 μm.

(C-D) Super-resolution confocal microscopy images show ventral optical sections from MDA-MB-231 cells, which were plated on gelatin (C) or collagen (D) and treated with EGF for 4 h prior to staining with phalloidin (F-actin) and antibodies against SEPT2, SEPT9_i1 and nucleoporins (NUP). Outlined regions (dashed rectangle) show in higher magnification septin localization at the nuclear rim (i) and across the ventral surface of the nucleus (ii). Overlaying regions between inverted magenta and green colors are shown in black. Scale bars, 10 μm.

(E) Super-resolution confocal microscopy image shows an optical mid-section from an MDA-MB-231 cell, which was plated on gelatin and treated with EGF for 4 h prior to staining with antibodies against SEPT9_i1, SEPT2 and nucleoporins (NUP). Outlined region is shown in higher magnification. Arrowheads point to septin fibers that localize to the nuclear envelope. Overlaying regions between inverted magenta and green colors are shown in black. Scale bar, 10 μm.

(F) Super-resolution confocal microscopy image of an apical optical section from an MDA-MB-231 cell, which was plated on gelatin and treated with EGF for 4 h prior to staining with phalloidin and antibodies against SEPT9_i1, SEPT7 and nucleoporins. Outlined region is shown in higher magnification. Arrowheads point to apical septin filaments that colocalize with F-actin. Overlaying regions between inverted magenta and green colors are shown in black. Scale bar, 10 μm.

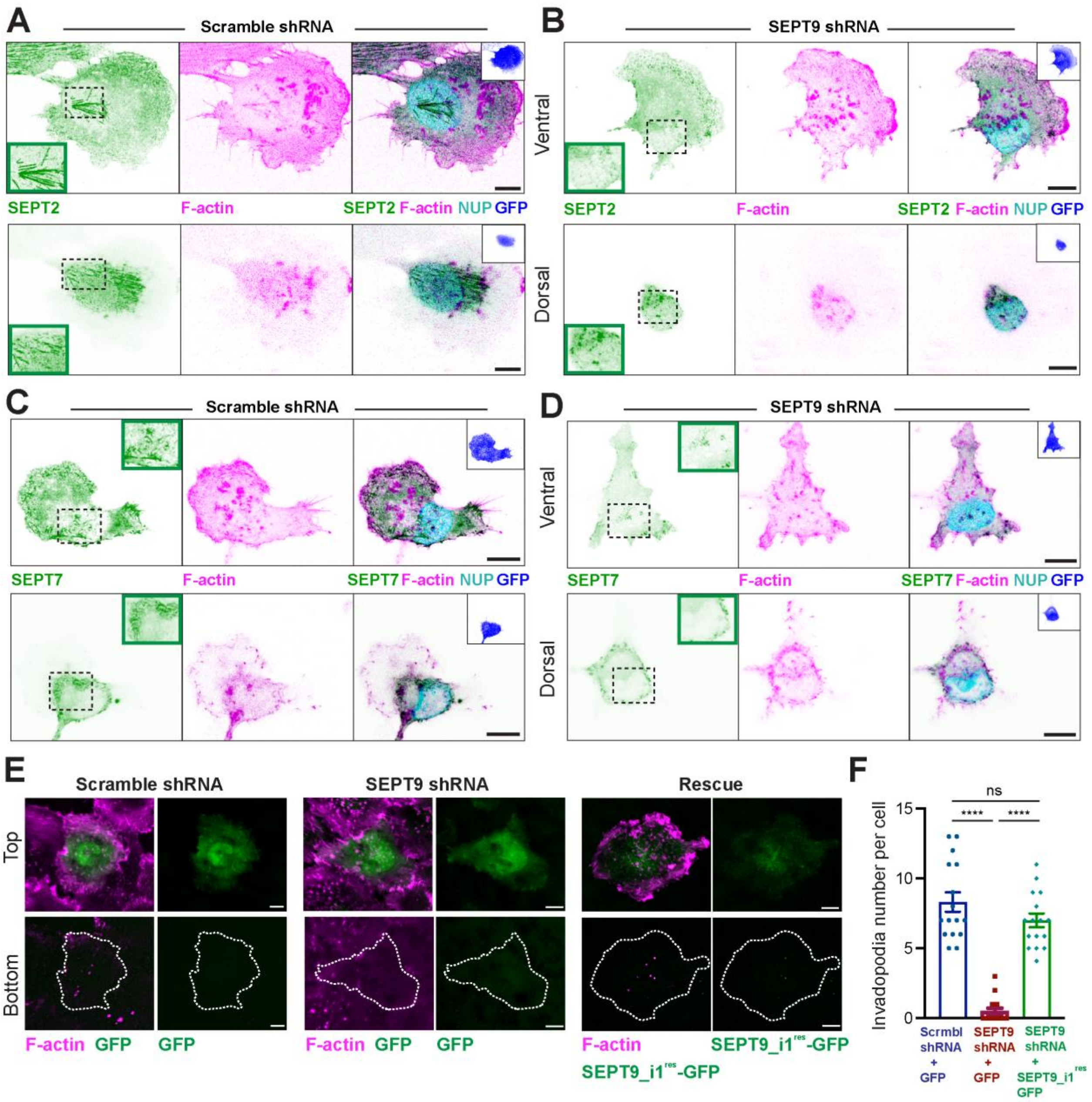


**Figure S2. Effects of SEPT9 depletion on SEPT2 and SEPT7 localization, and rescue of invadopodia phenotype, related to Figure 2.**

(A-B) Maximum intensity projections of ventral (top panels) and dorsal (bottom panels) sections acquired with confocal microscopy of MDA-MB-231 cells, which were treated with scramble control and SEPT9 shRNAs for 48 h. MDA-MB-231 cells were plated on gelatin and treated with EGF for 4 h prior to staining with phalloidin (F-actin; inverted magenta) and antibodies against SEPT2 (inverted green) and nucleoporins (NUP; inverted cyan). In SEPT2 panels, lower left inset shows in higher magnification the region outlined with dashed rectangle. Upper right inset shows the GFP fluorescence (inverted blue), which marks the expression of shRNAs. Scale bars, 10 μm.

(C-D) Maximum intensity projections of ventral (top panels) and dorsal (bottom panels) sections acquired with confocal microscopy of MDA-MB-231 cells, which were treated with scramble and SEPT9 shRNAs for 48 h. MDA-MB-231 cells were plated on gelatin and treated with EGF for 4 h prior to staining with phalloidin (F-actin; inverted magenta) and antibodies against SEPT7 (inverted green) and nucleoporins (NUP; inverted cyan). In SEPT7 panels, upper left inset shows the region outlined with a dashed rectangle in higher magnification. In the merge panels, upper right inset shows the GFP fluorescence (inverted blue), which marks the expression of shRNAs. Scale bars, 10 μm.

(E) Images show maximum intensity projections of confocal microscopy images, which were taken above (top) and under (bottom) a Matrigel-/laminin-coated Transwell membrane with MDA-MB-231 cells. Cells were transfected for 48 h with plasmids expressing GFP or SEPT9_i1$^{res}$-GFP (green), which was resistant to SEPT9 shRNA, and scramble or SEPT9 shRNA. Subsequently, cells were plated on Transwell membranes for 24 h in the presence of apicobasal EGF gradient, and stained with phalloidin (F-actin; magenta). Dashed lines outline the periphery of the cells above the basal side of the Transwell membrane, which contains the actin-rich protruding invadopodia. Scale bars, 10 μm.

(F) Quantification of the number invadopodia per cell (n = 16) that protrude into the basal side of the Transwell membrane. Data were statistically analyzed with a non-parametric one-way ANOVA Kruskal-Wallis test with Dunn's post-hoc analysis for pairwise comparisons.

n.s., not significant; ****, $p < 0.0001$

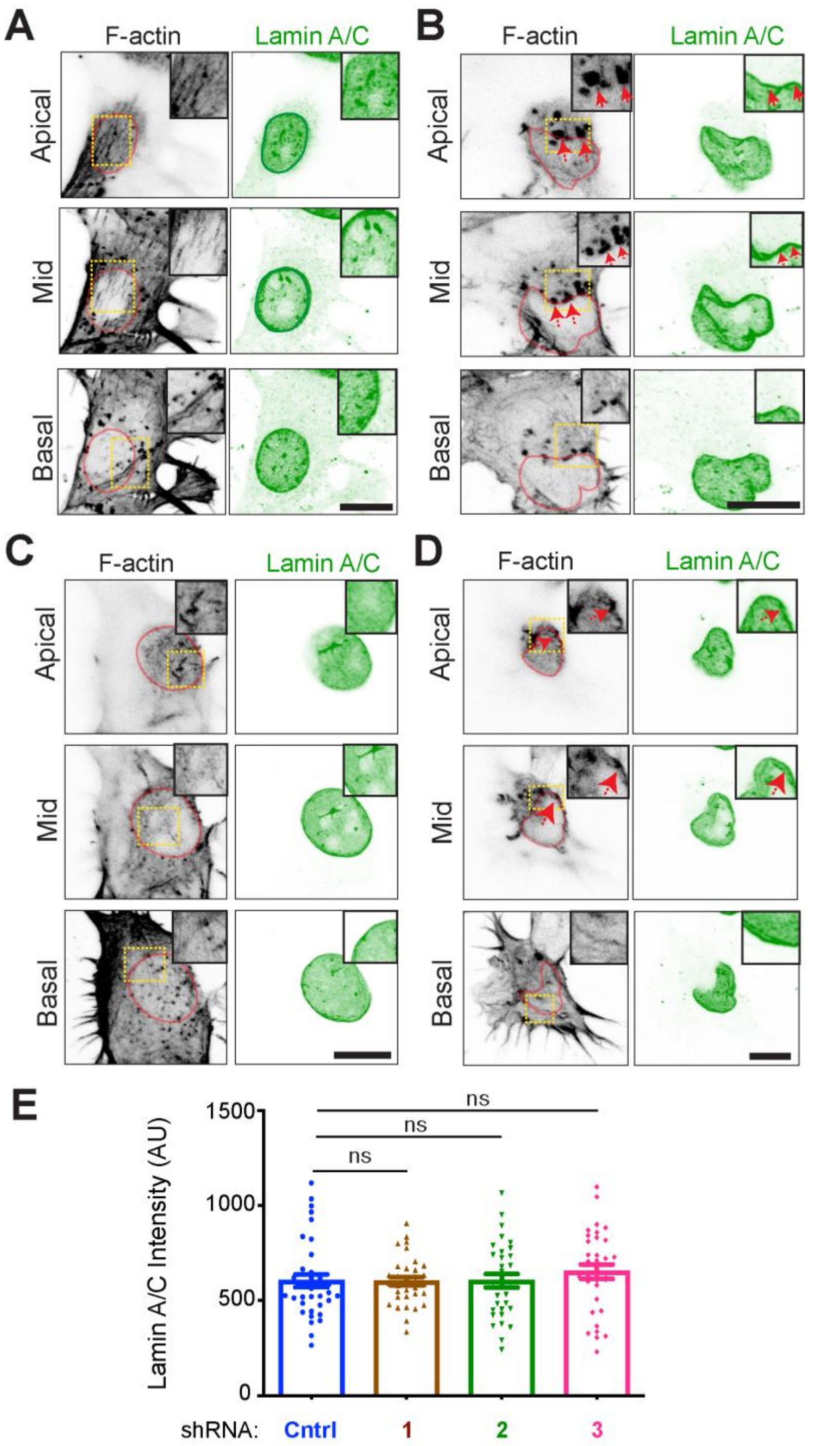


**Figure S3. SEPT9_i1 depletion disrupts actin organization but has no effect on lamin A/C expression, related to Figure 5.**

(A-D) Images show basal, medial and apical sections of MDA-MB-231 cells stained with phalloidin (F-actin) and lamin A/C antibody. Prior to imaging, MDA-MB-231 cells were transduced with lentiviruses carrying control shRNA (A) and shRNAs targeting SEPT9_i1 (shRNA1; B), SEPT9_i2 (shRNA2; C) or all SEPT9 isoforms (shRNA3; D), and were plated on gelatin for 5 h in the presence of EGF (10 nM for 4h). Nuclear footprints are outlined in red (red) based on the image of lamin A/C (green). Insets show in higher magnification the areas outlined by yellow dashed rectangles. Red arrows point to F-actin patches at regions of the nuclear rim. Scale bars, 10 μm.

(E) Bar graph shows quantification of endogenous lamin A/C levels in MDA-MB-231 cells (n = 32-36) which were transduced with lentiviruses carrying control shRNA (A) and shRNAs targeting SEPT9_i1 (shRNA1), SEPT9_i2 (shRNA2) or all SEPT9 isoforms (shRNA3), and were plated on gelatin for 5 h in the presence of EGF (10 nM for 4h). Cells were stained with antibodies to SEPT9_i1 and SEPT9_i2 to assess SEPT9 depletion. Representative images are shown in Figure 5A. Each data point is the average of the mean fluorescence of lamin A/C, which was derived from all optical sections of a single nucleus. Data were analyzed with a non-parametric one-way ANOVA Kruskal-Wallis test with Dunn's post-hoc analysis for pairwise comparisons. n.s., not significant

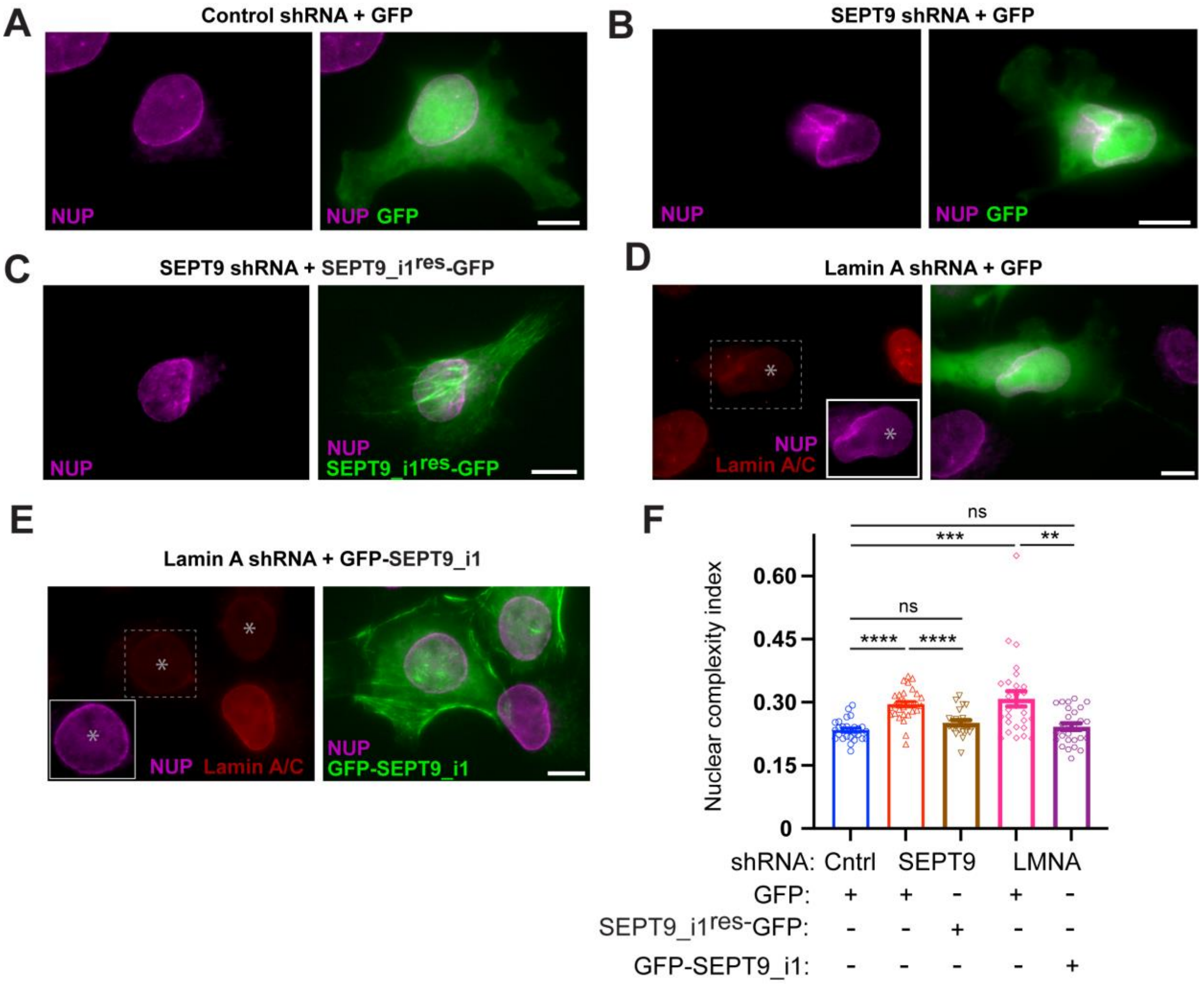


**Figure S4. SEPT9_i1 rescues the nuclear complexity phenotype of SEPT9 and lamin A knock-down, related to Figure 5.**

(A-E) MDA-MB-231 cells were transfected for 48-72 h with plasmids that express GFP (A-B, D) or GFP-tagged SEPT9_i1$^{res}$ (C, E), which is resistant to SEPT9 shRNA, and a non-targeting scramble control shRNA (A), SEPT9 shRNA (B, C) or lamin A shRNA (D, E). After plating on gelatin for 5 h in the presence of EGF (10 nM for 4h), cells were stained with antibody against nucleoporins (A-E) and lamin A/C (D-E). Images are maximum intensity projections of optical sections taken from the ventral and medial regions of cells. Asterisks mark nuclei with reduced lamin A/C levels due to lamin A shRNAs which are co-expressed with GFP (D) or GFP-SEPT9_i1$^{res}$ (E). Insets (D-E) show the nucleoporin stain of nuclei outlined in dashed rectangles. Scale bars, 10 μm.

(F) Quantification of the nuclear complexity index in MDA-MB-231 cells transfected with plasmids that express control scramble shRNA and GFP (n = 27), SEPT9 shRNA and GFP (n = 30), SEPT9 shRNA and SEPT9 shRNA resistant SEPT9_i1$^{res}$-GFP (n = 23), lamin A shRNA and GFP (n = 27) and lamin A shRNA and GFP-SEPT9_i1 (n = 28). Data were analyzed with a Welch ANOVA test with a post-hoc Dunnett's T3 test for multiple pair-wise comparisons.

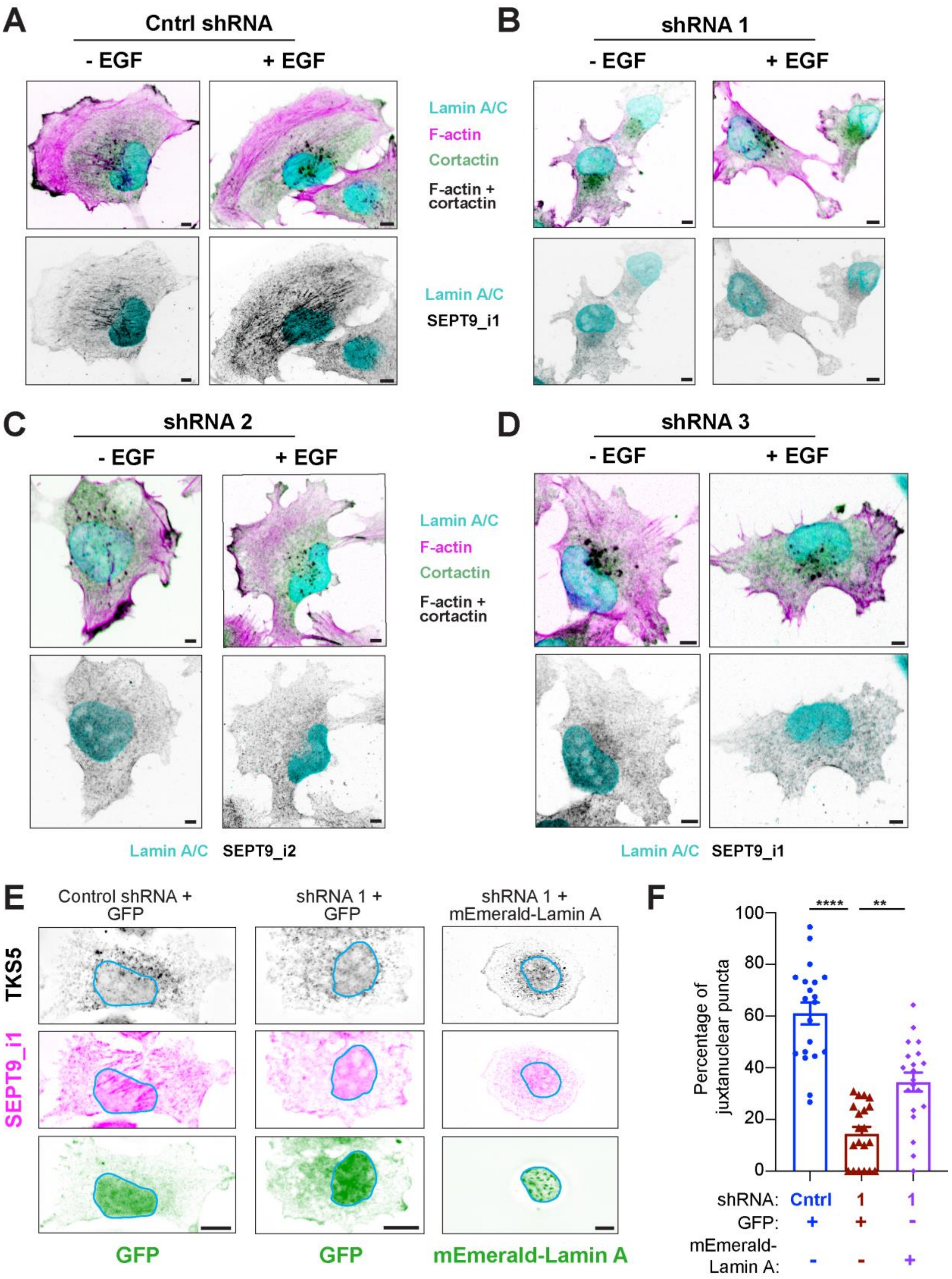

A
Cntrl shRNA
- EGF
+ EGF
B
shRNA 1
- EGF
+ EGF
Lamin A/C
F-actin
Cortactin
F-actin + cortactin
Lamin A/C
SEPT9_i1
C
shRNA 2
- EGF
+ EGF
D
shRNA 3
- EGF
+ EGF
Lamin A/C
F-actin
Cortactin
F-actin + cortactin
Lamin A/C SEPT9_i2
Lamin A/C SEPT9_i1
E
Control shRNA + GFP
shRNA 1 + GFP
shRNA 1 + mEmerald-Lamin A
TKS5
SEPT9_i1
GFP
GFP
mEmerald-Lamin A
F
****
**
Percentage of juxtanuclear puncta
100
80
60
40
20
0
shRNA: Cntrl 1 1
GFP: + + -
mEmerald-Lamin A: - - +

**Figure S5. SEPT9_i1 promotes EGF-induced enhancement of juxtanuclear invadopodia, related to Figure 6.**

(A-D) Images show ventral puncta with cortactin (inverted green) and actin (inverted magenta) with respect to the nucleus (lamin A/C; inverted cyan) in MDA-MB-231 cells, which were stained with antibodies to SEPT9_i1 or SEPT9_i2 (inverted grayscale). Prior to staining, MDA-MB-231 cells were treated with control shRNA (A) and shRNAs against SEPT9_i1 (shRNA1; B), SEPT9_i2 (shRNA2; C) and all SEPT9 isoforms (shRNA3; D) for 48 h and incubated with or without EGF (10 nM) for 4 h. Scale bars, 5 μm.

(E-F) Images (E) show maximum intensity of ventral optical sections of 786-0 cells stained for TKS5 (inverted monochrome) and SEPT9_i1 (inverted magenta). Prior to staining with antibodies, 768-0 cells were transduced with viruses carrying control or SEPT9 shRNAs for 48 h, and subsequently transfected with GFP or mEmerald-lamin A (inverted green) for 48 h prior to plating on gelatin for 5 h with EGF. Nuclear areas are outlined with blue lines. Images were processed with nearest neighbor deconvolution; white halo around the mEmerald-Lamin A fluorescence is due to out of focus haze, which was also present in the unprocessed image. Bar graphs (F) show the percentage of total ventral TKS5 puncta that localize within the nuclear footprint and a 3 μm-wide zone. Data (n = 20 cells per condition) were statistically analyzed with a Kruskal Wallis ANOVA test with a post-hoc Dunn's test for multiple pair-wise comparisons. **, $p < 0.01$; ****, $p < 0.0001$

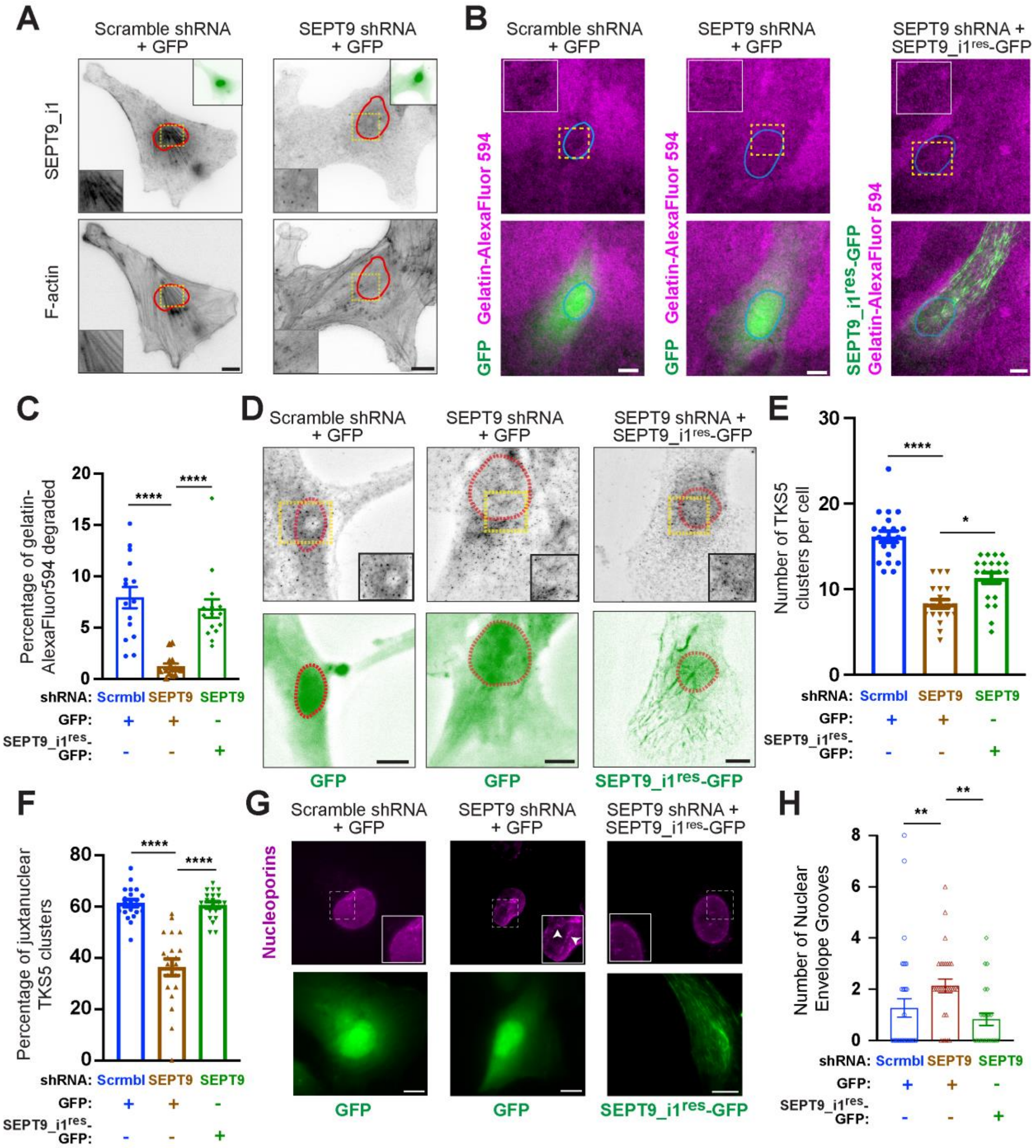


**Figure S6. SEPT9_i1 promotes the formation of juxtanuclear invadopodia and nuclear envelope integrity in 786-0 renal carcinoma cells, related to Figures 2, 3, 5 and 6.**

(A) Maximum intensity projections of ventral optical slices of 786-0 cells, which were transfected for 48 with plasmids that express GFP (insets, top right) and scramble control or SEPT9 shRNAs, and plated on gelatin for 5 h in the presence of EGF. Cells were stained with anti-SEPT9_i1 and phalloidin. Red lines outline nuclear areas, and insets (bottom left) show in higher magnification the areas in yellow dashed rectangles. Scale bars, 10 $\mu$m.

(B) Images show the gelatin-AlexaFluore594 matrix (magenta) underlying 786-0 cells, which were plated for 5 h in the presence of EGF after a 48 h transfection with plasmids expressing scramble control SEPT9 shRNA and GFP or shRNA-resistant SEPT9_i1$^{res}$-GFP(green). Areas of ECM degradation are shown as dark spots in images of gelatin-AlexaFluore594. Blue lines outline the areas occupied by nuclei, and regions in dashed rectangles are shown in higher magnification (insets). Scale bars, 10 $\mu$m.

(C) Quantification of the degraded gelatin-AlexaFluor594 that overlaps with the ventral nuclear footprint as percentage of the total surface area of the nucleus (n = 15 cells). Data were analyzed with a Kruskal Wallis ANOVA test with a Dunn's post-hoc multiple comparisons test.

(D) Images show maximum projections of ventral optical slices of TKS5 (inverted monochrome) stained 786-0 cells, which were plated on gelatin for 5 h in the presence of EGF after 48 h transfection with plasmids encoding for scramble control shRNA or SEPT9 shRNA and GFP or shRNA-resistant SEPT9_i1$^{res}$-GFP (inverted green). Nuclear areas are outlined with dashed red lines, and insets (lower right) show in higher magnification the areas outlined by the yellow dashed rectangles. Images were processed with nearest neighbors deconvolution. Scale bars, 10 $\mu$m.

(E-F) Bar graphs show the number of TKS5 clusters per cell (E) and percentage of total TKS5 clusters that overlap with the ventral nuclear footprint (F). Data (n = 20 cells per condition) were statistically analyzed with Kruskal-Wallis ANOVA test with a post-hoc Dunnett's T3 test for multiple pair-wise comparisons.

(G) Images show maximum intensity projections of nucleoporin-stained (magenta) 786-0 cells, which were plated on gelatin for 5 h in the presence of EGF after 48 h transfection with plasmids encoding for scramble control shRNA or SEPT9 shRNA and GFP or shRNA-resistant SEPT9_i1$^{res}$-GFP (green). Insets show outlined areas in higher magnification and arrowheads point to nuclear envelope grooves. Scale bars, 10 $\mu$m.

(H) Bar graphs show the number of nuclear envelope grooves quantified for cells transfected with scramble control shRNA and GFP (n = 33), SEPT9 shRNA and GFP (n = 29), and SEPT9 shRNA and SEPT9_i1$^{res}$-GFP (n = 24). Data were statistically analyzed with Kruskal-Wallis ANOVA test with a post-hoc Dunn's test for multiple pair-wise comparisons.

*, $p < 0.05$; **, $p < 0.01$; ****, $p < 0.0001$
